# One Model to Rule Them All: An Integrative Approach to Matrix-Based Analyses in Neuroimaging Connectomics

**DOI:** 10.1101/459545

**Authors:** Gang Chen, Paul-Christian Bürkner, Paul A. Taylor, Zhihao Li, Lijun Yin, Daniel R. Glen, Joshua Kinniso, Robert W. Cox, Luiz Pessoa

## Abstract

Network modeling in neuroimaging holds promise in probing the interrelationships among brain regions and potential clinical applications. Two types of matrix-based analysis (MBA) are usually seen in neuroimaging connectomics: one is the functional attribute matrix (FAM) of, for example, correlations, that measures the similarity of BOLD response patterns among a list of predefined regions of interest (ROIs). Another type of MBA involves the structural attribute matrix (SAM), e.g., describing the properties of white matter between any pair of gray-matter regions such as fractional anisotropy, mean diffusivity, axial and radial diffusivity. There are different methods that have been developed or adopted to summarize such matrices across subjects, including general linear models (GLMs) and various versions of graph theoretic analysis. We argue that these types of modeling strategies tend to be “inefficient” in statistical inferences and have many pitfalls, such as having strong dependence on arbitrary thresholding under conventional statistical frameworks.

Here we offer an alternative approach that integrates the analyses of all the regions, region pairs (RPs) and subjects into one framework, called Bayesian multilevel (BML) modeling. In this approach, the intricate relationships across regions as well as across RPs are quantitatively characterized. This integrative approach avoids the multiple testing issue that typically plagues the conventional statistical analysis in neuroimaging, and it provides a principled way to quantify both the effect and its uncertainty at each region as well as for each RP. As a result, a unique feature of BML is that the effect at each region and the corresponding uncertainty can be estimated, revealing the relative strength or importance of each region; in addition, the effect at each RP is obtained along with its uncertainty as statistical evidence. Most importantly, the BML approach can be scrutinized for consistency through validation and comparisons with alternative assumptions or models. We demonstrate the BML methodology with a real dataset with 16 ROIs from 41 subjects, and compare it to the conventional GLM approach in terms of model efficiency, performance and inferences. Furthermore, we emphasize the notion of full results reporting through “highlighting,” instead of through the common practice of “hiding.” The associated program will be available as part of the AFNI suite for general use.

## Introduction

Conventional neuroimaging data analyses adopt a massively univariate approach through voxel-wise modeling, and also require an additional step of correction for multiple testing due to the inefficient modeling approach through the pretense that each spatial unit is unrelated to its neighbors (Chen et al., 2018a). Although relatively robust, such an approach only allows one to reach conclusions about localized effects. As our knowledge regarding information processing accumulates, it has become more appealing to explore the interactions among brain regions, e.g., by network modeling to explore how information is shared and related across regions. It is believed that differences in information flow within a network may exist between various patient groups and controls, holding the promise for potential clinical applications on psychiatric and neurological disorders (e.g, Alzheimer’s: Chong et al., 2017; autism and schizophrenia: Mastrovito et al., 2018).

Network modeling usually reduces the dimensionality of the whole-brain time series data to a matrix format that symbolizes the relationships among a set of brain regions. For example, with *m* regions of interest (ROIs) defined across all subjects, the investigator summarizes the original data into a *m* × *m* matrix for each subject. Two broad categories of matrix-based analysis (MBA) exist in neuroimaging connectomics, depending on whether the study is functional or structural. For the former, a researcher could define a functional attribute matrix (FAM) using the inter-region correlations^1^ (IRCs) of the BOLD signal among the ROIs. For the latter, one can generate a structural attribute matrix (SAM) that typically summarizes properties of white matter (WM) connections between any pair of gray matter ROIs; for example, in a DTI study, one might perform tractography to estimate the locations of WM between pairs of gray matter targets and then analyze quantities such as average fractional anisotropy (FA), mean diffusivity (MD), axial diffusivity (AD or L1), etc. of each estimated connection (e.g., Taylor et al. 2016). Without loss of generality, we focus our discussion here mainly on the case of IRC matrices, which can be readily extended to the SAM analysis.

With one group of *n* subjects and *m* > 2 regions, *R*_1_,*R*_2_,.…, *R_m_*, the total number of unique effect estimates is 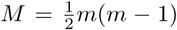. For the *k*th subject (*k* = 1, 2,…, *n*), the estimated values (e.g., correlation coefficients {*r_ijk_*, *i* > *j*}) correspond to *M* region pairs, and they form a symmetric (*r_ijk_* = *r_jik_*, *i*, *j* = 1, 2,…, *m*) *m* × *m* positive semi-definite matrix 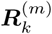 with diagonals *r_iik_* = 1 (Fig. 1, left). In the case of correlation coefficients, their Fisher transformed version 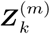 (Fig. 1, right) through *z* = *arctanh*(*r*) is usually adopted during analysis so that methods assuming Gaussian distribution may be utilized as Fisher *z*-values are more likely to be Gaussian-distributed than raw Pearson correlation coefficients. The research of interest herein is focused on the population average effect of each region pair (*i*,*j*). Because 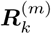 and 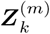 are both symmetric in (*i*,*j*), inferences at the population level can be made through the *M* elements in the lower triangular part (shaded gray in Fig. 1).

**Figure 1:**
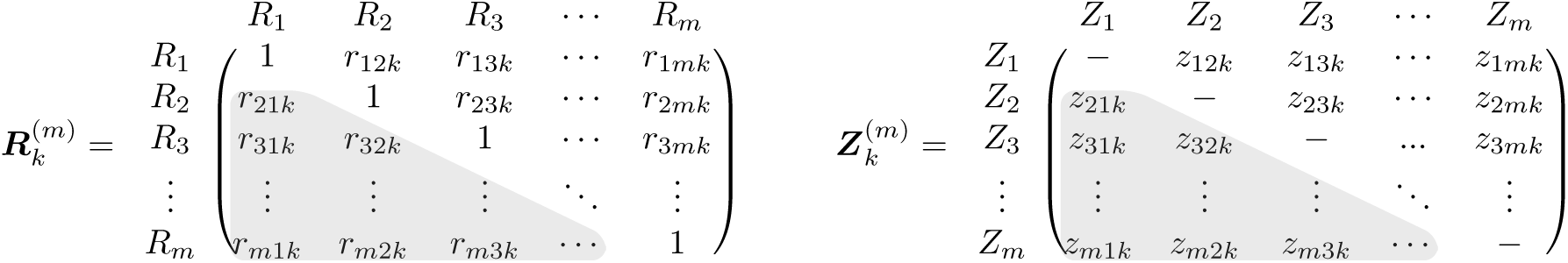
Inter-region correlation (IRC) matrix 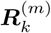 among the *m* regions for the *k*th subject and its Fisher-transformed counterpart 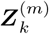. Due to the symmetry, only half of the off-diagonal elements (shaded in gray) are usually considered during IRC analysis.

The general interest of MBA at the population level is the statistical inference about the population effect for each RP. However, a complex issue to manage is that each matrix element is correlated with some of others, similar to the situation with inter-subject correlation (ISC) data structure (Chen et al., 2017). Suppose that *z*_*i*_1_*j*_1_*k*_ and *z*_*i*_2_*j*_2_*k*_ are two *z*-values that are associated with the IRCs of the *k*th subject, *r*_*i*_1_*j*_1_*k*_ and *r*_*i*_2_*j*_2_*k*_, of two region pairs (RPs). We denote the correlation between any two elements, *z*_*i*_1_*j*_1_*k*_ and *z*_*i*_2_*j*_2_*k*_, that pivot around a common region (e.g., *i*_1_ = *i*_2_) as *ρ*, with the assumption that the relatedness *ρ* remains the same across all regions. In other words, *ρ* characterizes the interrelatedness of *z*_i_1_*j*_1_*k*_ and *z*_*i*_1_*j*_2_*k*_ among the three regions among which the two RPs share a common region. To consider the group-wide set of IRCs, we further define ***z***_*k*_ = *vec*({*z_ijk_*, *i* > *j*}) to be the vector of length *M* whose elements are the column-stacking of the lower triangular part of the matrix ***Z***^(*m*)^ in Fig. 1. That is, ***z*** is the half-vectorization of 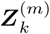 excluding the main (or principal) diagonal: 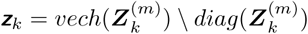. The variance-covariance matrix of ***z***_*k*_ can be expressed as the *M* × *M* matrix,

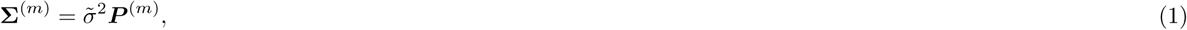

where 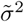 is the variance of *z_ijk_*, *i* > *j*, and ***P***^(*m*)^ is the correlation matrix that is composed of 1 (diagonals), *ρ* and 0. An example of ***P***^(5)^ is shown in Fig. 2. It has been analytically shown (Chen et al., 2016) that —1/[2(*m* — 2)] ≤ *ρ* ≤ 0. 5 (when *m* > 3) at the individual subject level, and because of the presence of correlations among some elements of 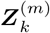, it becomes crucial to capture this correlation structure ***P***^(*m*)^in any modeling framework.

**Figure 2:**
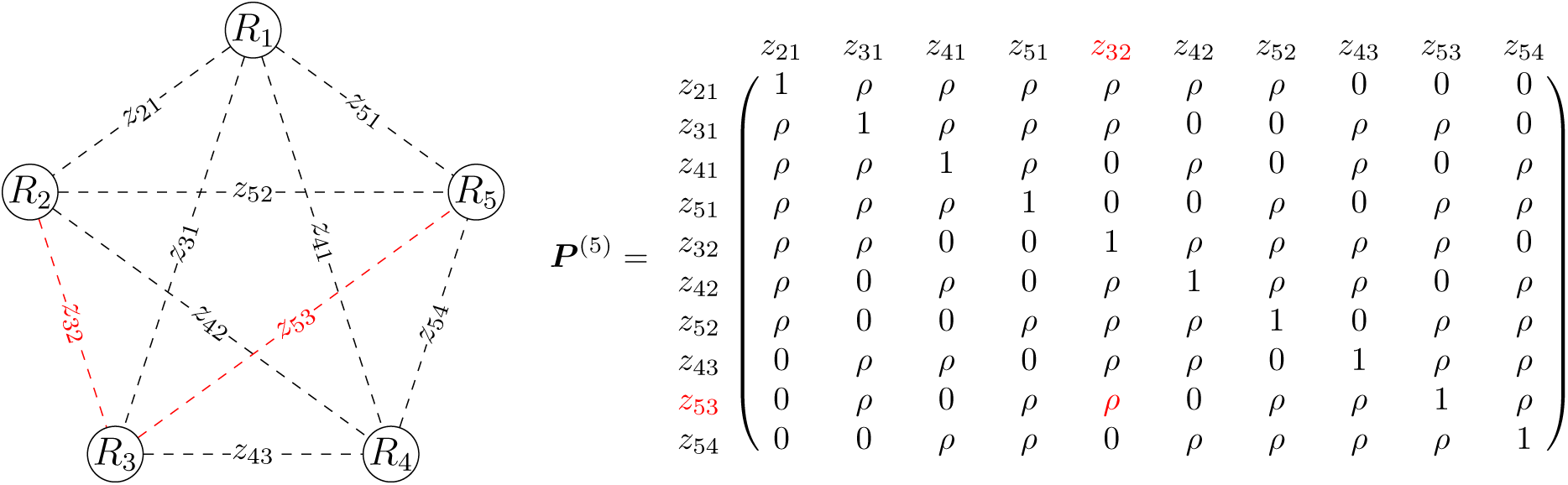
IRC with *m* = 5 regions. Left: pictorial representation of 5 × 5 region pairing. Unlike the typical representations with solid lines in the field, we intentionally adopt dashed lines here to indicate correlations, not real connections, between regions to avoid potential misinterpretations. Right: The complex relatedness among the off-diagonal elements in 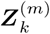 is illustrated with the correlation matrix ***P***^(5)^ for *m* = 5 regions, in which *ρ* represents the correlation when two elements (e.g., *z*_32_ and *z*_53_, colored in red) are associated with a common region (e.g., *R*_3_). Without loss of generality, the third index *k* in *z_ijk_* for subjects is dropped here for clarity.

In the case of white matter analysis (e.g., using diffusion-based tractography), the measure matrix is usually not full in the sense that direct connections are not observed between some RPs. That is, for a given subject some elements in the data matrix 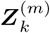 are missing or zero, the number of RPs in the input data is usually less than *M*, and the missingness may not be necessarily the same across subjects (e.g., Taylor et al., 2016). However, for convenience in the present description, we focus on the case of IRC, which usually does not have missing elements. We will elaborate on the missingness issue in the Discussion section and extend the modeling strategy to SAM of the DTI case.

## MBA with conventional approaches

An intuitive and straightforward approach to making generalization at the population level is to focus on each RP and formulate a general linear model (GLM) for the *l*th RP (*l* = 1, 2,…, *M*),

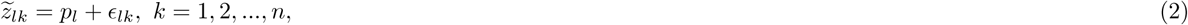

where 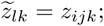 *l* indices the flattened list of RPs (*i*, *j*),*i*, *j* = 1, 2, …,*m* (*i* > *j*); *pl* represents the population effect of the *l*th RP, and *∊_lk_* is the deviation of the *k*th subject on the *l*th RP, which is assumed to follow a Gaussian distribution. Each of the *M* models in (2) is essentially a Student’s *t*-test for the null hypothesis of *H*_0_: *p_l_* = 0. This modeling strategy has been incorporated into neuroimaging tools such as network-based statistics (NBS) implemented in Matlab (Zalesky et al., 2010) and FSLNets in FSL (Jenkinson et al., 2012).

For the convenience of model comparisons, the M separate models in (2) can be merged into one GLM by pooling all the residuals across the *M* RPs (e.g., through treating the RPs as *M* levels of a factor in the model),

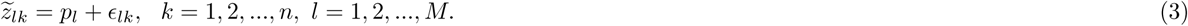

Furthermore, the GLM platforms (2) and (3) can be readily extended to cases with categorical variables (e.g., between-subjects factors such as two or more groups, or within-subject factors such as conditions or tasks) or with quantitative explanatory variables (e.g., subject-specific values such as age and behavioral measures).

The immediate challenges that the modeling frameworks in (2) and (3) face are twofold. First, the relationships among the *M* RPs are implicitly involved to some extent in the process of correction for multiple testing, such as in the permutation-based methods in NBS and FSL Randomise; however, those intricate relationships, as illustrated in the seemingly irregular pattern of the correlation matrix ***P***^(*m*)^ for those *M* elements, are not accurately quantified under GLM, at least not explicitly characterized to the extent of the accurate pattern as shown in ***P***^(5)^ of Fig. 2. The second challenge is the issue of multiplicity: as there are a total of *M* models under the GLM framework (2) that correspond to the *M* RPs, it remains a daunting job to effectively and efficiently maintain an overall false positive rate (PFR) under the null hypothesis significance testing (NHST) framework, as evidenced by the variety and breadth of current correction methods (Zalesky et al., 2010; Baggio et al., 2018).

Another presently popular approach for analyzing subject matrices at the group level is graph theoretic analysis. Typically a hard threshold is chosen at some stage either for the elements in the correlation matrix (e.g., 0, 0.2, 0.25, etc.) or for the percentile of the “surviving paths” (e.g., 15%) at the individual subject level. Thereafter multiple steps lead to a long list of topological features that are extracted at micro-, meso- and macro-scales. It remains unclear as to how these thresholds translate to physiological features at the neurological level, and the robustness of the final results is questionable after a cascade of summarization steps, each of which involves thresholding or ignores uncertainties from the previous steps (e.g., see the issue of garden of forking paths, Gelman and Loken, 2013; Chen et al., 2018).

This rest of the paper is structured as follows. First we formulate a population analysis through an ANOVA or linear mixed-effects (LME) platform by pivoting the ROIs as the levels of a random-effects factor, and we then convert the LME model to its associated Bayesian multi-level (BML) counterpart. We describe how the BML weighs and pools the information based on the precision across all the regions and RPs. As a practical example, we apply the modeling approach to an experimental dataset of a 16 × 16 correlation matrix from each of 41 subjects. In the Discussion, we review current MBA methodologies in neuroimaging, and elaborate the advantages and limitations of BML modeling for MBA. Major acronyms and terms are listed in Table 1.

**Table 1:** Acronyms and terminology.

|  |  |  |  |
| --- | --- | --- | --- |
| BML | Bayesian multilevel | LOO-CV | leave-one-out cross-validation |
| ESS | effective sample size | LOOIC | leave-one-out information criterion |
| FAM | functional attribute matrix | MBA | matrix-based analysis |
| FPR | false positive rate | MCMC | Markov chain Monte Carlo |
| GLM | general linear model | NHST | null hypothesis significance testing |
| HMC | Hamiltonian Monte Carlo | PPC | posterior predictive check |
| ICC | interclass correlation | ROI | region of interest |
| IRC | inter-region correlation | RP | region pair |
| LME | linear mixed-effects | SAM | structural attribute matrix |

## Theory: MBA through Bayesian multilevel modeling

Throughout this article, italic letters in lower case (e.g., α) stand for scalars and random variables; lowercase, boldfaced italic letters (***α***) for column vectors; Roman and Greek letters, respectively, for fixed and random effects in the conventional statistics context such as ANOVA and LME on the righthand side of a model equation. Although the terms of “fixed” and “random” effects are genuinely non-Bayesian, we still use them here as we expect most readers to be familiar with the conventional terminology. For instance, a conventional fixed-effects parameter under ANOVA and LME is treated as constant that is shared by all entities (e.g, subjects, ROIs), and a random-effect parameter as variable because it differs from one entity (e.g., subject, ROI) to another. The conventional distinction of fixed- vs. random-effects is replaced by one that separates the modeling decision (a parameter as varying or non-varying) under the Bayesian framework from the inference decision (e.g., prior choices or partial pooling) (Gelman, 2005).

### Bayesian modeling based on three-way random-effects ANOVA

We start with a framework of MBA for *m* ROIs in the brain through a three-way random-effects ANOVA or LME with three crossed random-effects components,

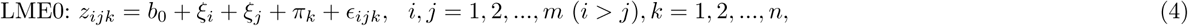

where *b*_0_ is the intercept or overall effect that is shared by all regions, RPs and subjects; *ξ_i_* and *ξ_j_* are the random effects or deviations from the population effect *b*_0_ for the two ROIs *i* and *j*, respectively; *π_k_* is the random effect attributable to subject *k*; and *∊_ijk_* is the residual term. Due to the symmetric nature of the data structure in 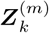, only half of the matrix elements excluding the diagonals (either the lower or upper triangular part of the matrix) are utilized in the model (4), and thus the index inequality of *i* > *j* is placed for the input data. The three random-effects components, *ξ_i_*, *ξ_j_* and *π_k_*, form a crossed (or cross-classified) structure with a factorial (or combinatorial) layout among the levels (or indices *i*, *j* and *k*) of the three factors.

With the assumption of independent Gaussian distributions, 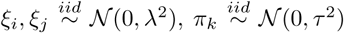, and *∊_ijk_* ~ 𝒩(0, σ^2^), the model in (4) can be solved under a three-way random-effects ANOVA or LME. Unlike the *M* separate GLMs in (2) or the pooled version (3) that treats the *M* RPs as separate and independent entities, each effect *z_ijk_* is decomposable as the additive effects of multiple components under the LME model (4). Such a decomposition, even though still vulnerable to assumption violations, allows for more accurate effect characterization and more powerful inferences than the typical GLM approach of analyzing each RP separately as in (2) and (3). For instance, related to the concept of intraclass correlation (ICC), the correlation between two RPs, (*i*, *j*_1_) and (*i*, *j*_2_) (*j*_1_ ≠ *j*_2_) can be derived as,

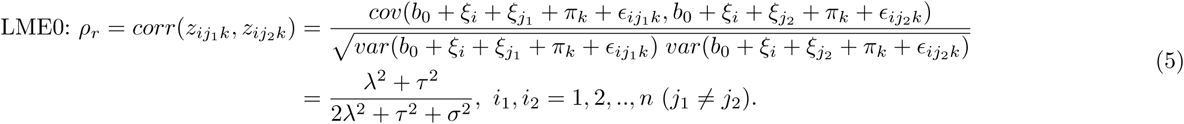

due to the fact that the two RPs share a common region *R_i_*. Unlike the GLM approaches such as (2) and (3) where the RPs are pretended to be unrelated during the analysis, the integrity of relatedness among the IRC matrix elements, as shown in ***P***^(*m*)^, is maintained under the LME0 model (4) as characterized in (5).

It is worth noting that the relatedness, as shown in (5), at the population level between two IRC elements that pivot at one common region, *ρ_r_*, is not constrained by the relationship of –1/[2(*m* — 2)] ≤ *ρ* ≤ 0.5 (*m* > 3) that was proved at the individual level (Chen et al., 2016). The reason is that there are three random-effects components involved at the group level, as specified in the model formulation (4), which are more than the number of components (i.e., two) at the individual level as well as the number of components with the inter-subject correlation scenarios (Chen et al., 2016).

Similarly, the correlation of an RP (*i*, *j*), between two subjects *k*_1_ and *k*_2_ can be derived as,

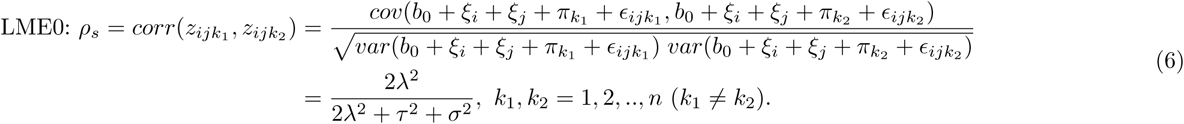

due to the fact that their effects are measured at the same RP.

However, there are two hurdles associated with the LME0 model (4) that have to be overcome. The first challenge is that, even though the random-effects components, *ξ_i_* and *ξ_j_*, that are associated with the two regions *i* and *j*, are assumed to follow the same Gaussian distribution 𝒩(0, λ^2^) (that is why we denote them by the same symbol ξ), they would have to be treated as two separate random-effect components in practice when one solves the system. Furthermore, due to the fact that only half of the off-diagonal elements in the matrix 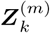 are utilized as input, the two random-effects components *ξ_i_* and *ξ_j_* are generally not evenly arranged among all the RPs, leading to unequal estimation of the two random-effects components. The problem can be resolved through using both upper and lower triangular off-diagonal elements of the matrix as input, as previously adopted in LME modeling for inter-subject correlation analysis (Chen et al., 2016). Therefore, the theory-based index constraint of *i* > *j* under the model LME0 in (4) is relaxed to *i* ≠ *j* for numerical implementations. The second hurdle is that, under the conventional modeling frameworks such as ANOVA and LME, we can only obtain the point estimate and the associated uncertainty (e.g., standard error) for each fixed effect (e.g, the overall effect *b*_0_ in the model LME0), as well as the variances for those random-effect components (e.g., λ^2^, τ^2^, and *σ^2^*). However, the typical research focus under the current context is to make inference about each RP, that is, about *p_ij_* = *b*_0_ + *ξ_i_* + *ξ_j_*, which cannot be achieved under the conventional modeling frameworks such as ANOVA and LME. In other words, the second hurdle basically renders the standard LME modeling framework infeasible for statistical inferences of MBA at the region and RP levels.

To be able to derive the effect at each region and RP directly, we reformulate the effect decomposition of *z_ijk_* under the BML framework, following the same modeling strategy in our previous work (Chen et al., 2018). For example, we translate the model LME0 in (4) to its BML counterpart^2^, forming a hierarchical or multilevel structure with data clustered by region, RP, and subject,

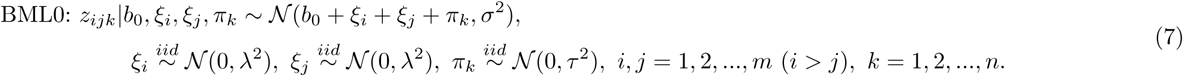

Both of the above hurdles under the LME model (4) are overcome now under the BML system (7). First, only half of the off-diagonal elements (e.g., the lower triangular part) in ***Z***^(*m*)^ are needed as input under BML through a numerical implementation of multi-membership modeling scheme^3^ (Bürkner, 2018). Second, with a prior (e.g., noninformative uniform distribution) for *b*_0_, the BML system (7) can be analyzed and the posterior distribution for each RP can be assembled through

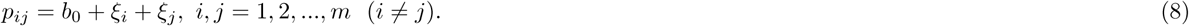

In addition, the effects that are attributable to each region, *r_i_*, and each subject, *s_k_*, as well as their interaction (a subject’s effect at a particular region), *t_ik_*, can be derived as well through their posterior distributions with

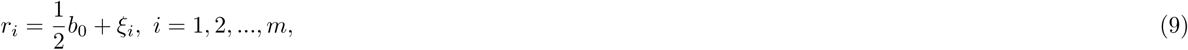

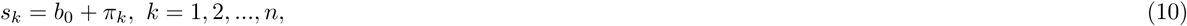

and

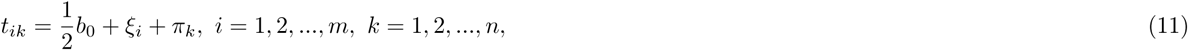

respectively. The intercept *b*_o_ is the overall effect shared by all regions, RPs and subjects, which may or may not be of interest to the investigator. The factor of 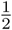 in the region-specific effect formula of *r_i_* (9) and in the region-subject interaction effect *t_ik_* (11) reflects the fact that the effect of each RP is evenly shared between the two associated regions. The region-specific effects can be utilized to assess the effect (contribution or importance) of an ROI relative to all other regions. Similarly, the effect of a subject can be used to judge whether the subject is an outlier relative to the whole group.

The BML framework (7) offers a good opportunity to discuss and substantiate the avoidance of the conventional terminology of “fixed- vs. random-effects.” Being of research interest for statistical inference, the effect at each region or each RP, on one hand, would be considered as “fixed” under the conventional framework; on the other hand, such an effect is modeled as random in the LME0 model (4). Such a conceptual inconsistency automatically dissolves once we abandon the distinction of fixed- vs. random-effects and instead differentiate two different types of effects: the effect *ξ_i_* associated with each region (or *p_ij_* in (8) associated with each RP) is modeled under the model BML0 (7) for the sake of statistical inference through partial pooling with a Gaussian prior, whereas the subject-specific effect *π_k_* in the BML framework (7) represents the varying component across subjects. In other words, the distinction of “fixed- vs. random-effects” under the conventional framework is mapped to the differentiation, in the current context, between information pooling across regions and cross-subject variability.

### Further extensions of BML for MBA

The LME0 model in (4) can be expanded or generalized by including two types of random-effects interaction components: one component is the RP-specific term (i.e., the interaction between two regions), and the other component is the interaction between a region and a subject. The expansions lead to three more LME models, corresponding to three different combinations of the two extra effects, as shown below,

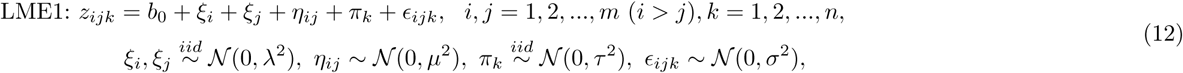

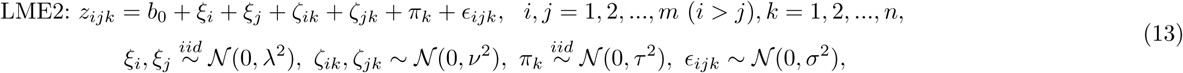

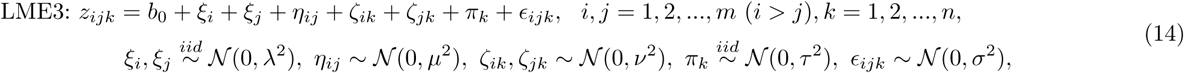

where η_*ij*_ is the effect of the RP that is associated with regions *i* and *j* (i.e., the interaction effect between regions *i* and *j*) relative to the overall effect *b*_0_ and the two region effects,*ξ_i_* and *ξ_j_*, while *ζ_ik_* and *ζ_jk_* are the interaction effects between region *i* and subject *k* as well as the interaction between region *j* and subject *k*, respectively. We note that the region-specific effect η*_ij_* captures the unique effect of each RP in addition to the overall effect *b*_0_ and the common effects from the two involved regions, *ξ_i_* and *ξ_j_*; the same subtlety applies to the region-subject interactions *ζ_ik_* and *ζ_jk_*.

The two ICC measures in (5) and (6) can be correspondingly updated to the following for the three LME models,

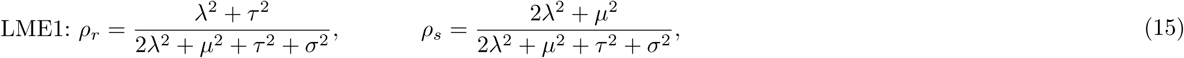

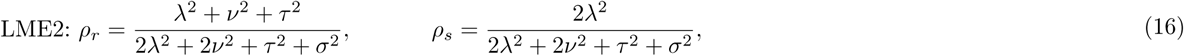

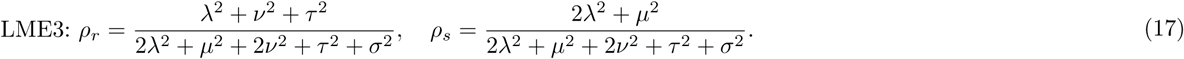

Among the four LME models, LME0 is the simplest and LME3 is the most complex and inclusive, while LME1 and LME2 are intermediate. As the number of components in a model increases, so does the number of parameters to be estimated. For example, with 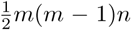 data points *z_ijk_* input, the total number of parameters involved at the
right-hand side of the model LME3 in (14) is *m*(*m* −1)+*mn*+2.For the model LME3 to be identifiable, it is a prerequisite that the following relationship be met,

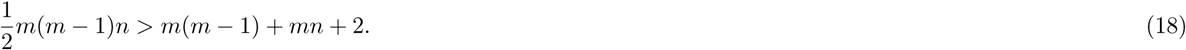

Traditionally, these LME models can be compared based on the tradeoff between model performance and complexity (e.g., number of parameters), with likelihood ratio testing and information criteria such as Akaike information criterion (AIC) and the so-called Bayesian information criterion (BIC) (Bates et al., 2015).

We further consider two types of BML extension based on the primary model BML0 in (7). The first type involves potential interaction effects, in parallel with the three LME expansions from LME0. Specifically, by incorporating the interaction effect between the two regions of each RP as well as the interaction effect between each region and each subject, we have three more BML models (corresponding to the LME models of the same index):

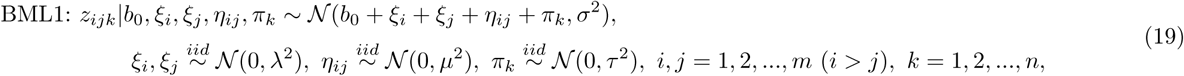

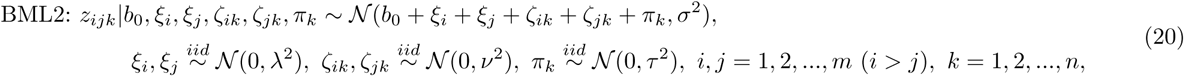

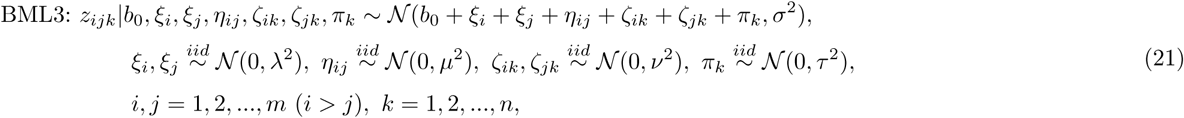

where η_*ij*_ is the RP-specific effect or the interaction between regions *i* and *j*, while *ζ_ik_* is the interaction effect between region *i* and subject *k* and *ζ_jk_*, between region *j* and subject *k*. The two interaction effects, *ζ_ik_* and *ζ_jk_*, are considered as two members, *i* and *j*, of a multi-membership cluster. Because of the sheer number of parameters, their LME counterparts are not always identified (e.g., because the prerequisite (18) is violated), but these BML models can be analyzed under the Bayesian scheme because of the constraints and regularization applied through priors. Similar to the LME case, complexity increases from BML0 to BML3.

Under the extended BML models, the region and RP effects can be similarly derived from their posterior distributions as for BML0 in (7). The region- and subject-specific effect formulas as well as their interactions remain the same for all these extended models as (9), (10), and (11), respectively. Along the same vein, the RP-specific effect formulation remains the same as (8) for BML2 (20), while for BML1 (19) and BML3 (21) it changes to

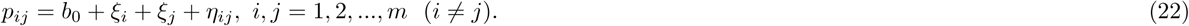

A natural question is, how to decide which model is most appropriate to use among the four BML variations, BML0, BML1, BML2, and BML3? In other words, which and how many interaction terms should one consider among the various choices? One important aspect of the Bayesian framework is model quality check through various prediction accuracy metrics. The aim of the quality check is not to accept or reject the model, but rather to check whether it fits the data well. For instance, posterior predictive check (PPC) simulates replicated data under the fitted model and then graphically compares actual data to the model prediction. The underlying rationale is that, through drawing from the posterior predictive distribution, a reasonable model should generate new data that look similar to the acquired data at hand.

One solution to model comparisons is to adopt Pareto smoothed importance-sampling leave-one-out cross-validation (LOO-CV) (Vehtari et al., 2017). This approach compares the potential model candidates by estimating the point-wise, out-of-sample prediction accuracy from each fitted Bayesian model using the log-likelihood evaluated at the posterior simulations of the parameter values, and selects the one with the lowest information criterion (if selection of a single model is desired). This accuracy tool uses probability integral transformation (PIT) checks, for example, through a quantile-quantile (Q-Q) plot to compare the LOO-PITs to the standard uniform or Gaussian distribution (Vehtari et al., 2017).

Another approach to comparing models is to visually inspect model prediction accuracies against the original data. For example, PPCs can be employed to graphically compare competing model to the actual data. Furthermore, as a model validation tool, PPC intuitively provides a visual tool to examine any systematic differences and potential misfit of the model, similar to the visual examination of plotting a fitted regression model against the original data. We demonstrate these model quality check techniques in the real data analysis example, below.

In addition, not only can model comparisons be performed among Bayesian candidates, but also can such comparisons be feasible between BML and GLM by fitting a GLM under a Bayesian framework. For example, the conventional GLM approach can be directly compared to the BML variations through cross validations (e.g., assessing posterior predictive accuracy via PPCs). Such comparisons are possible by assigning a noninformative prior to the model parameters as shown with, for example, the GLM formulation (3) Bayesianized to

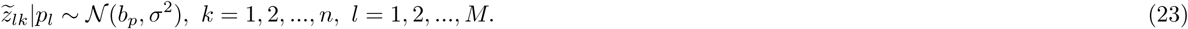

#### Incorporation of explanatory variables under BML for MBA

A second type of model extension is the incorporation of one or more subject-specific (e.g., sex, age, behavioral measures) or within-subject (e.g., multiple conditions such as positive, negative and neutral emotions) explanatory variables. For example, with one explanatory variable, BML1 in (19) can be directly augmented by adding a subject-level covariate *x_k_* to,

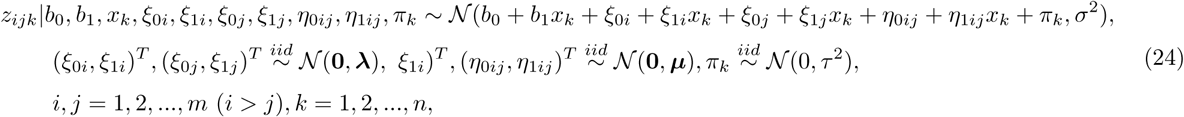

where **λ** and *μ* are 2 × 2 variance-covariance matrices. Model comparisons can also be performed among various candidate models at the presence of explanatory variables, with options similar to those suggested above.

In addition to the region- and subject-specific effects such as *r_i_* defined in (9) and *s_k_* in (10) that can be applied to the BML model (24), the region and subject interaction and the region- and RP-specific effects associated with the covariate *x* under the BML (24) can be similarly obtained through

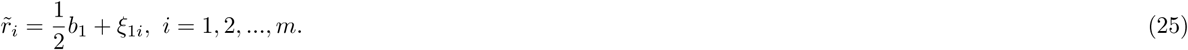

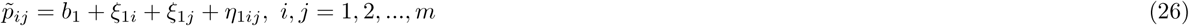

Lastly, the interaction effect between region and subject under the BML (24) is,

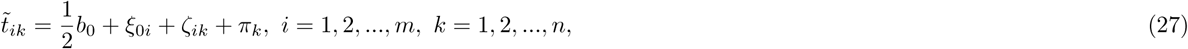

Cases with more than one explanatory variable can be similarly formulated as in the BML model (21) or (24). The research interest can be region-specific (e.g., *r_i_*) or RP-specific (e.g., *p_ij_*) effects for the BML models; alternatively, the research interest can be region-specific (e.g., *r̃_i_*) or RP-specific (e.g., *p̃_ij_*) effects of the explanatory variable *x*, for example, for the BML model (24). It is worth emphasizing that a unique feature of BML modeling is that the region (and subject) effect can be obtained through the posterior distribution of, for example, *ξ_i_* and *π_k_*, so that the investigator could 1) quantitatively derive the relative importance of each ROI within the model framework and naturally quantify the uncertainty for each effect of interest; and 2) investigate as to which subjects are more outlying than others, or explore the possibility of including potential covariates based on the outlying information. This is a major benefit over ANOVA and LME, because such inferences on the effects of each region and each RP cannot be achieved under the conventional frameworks.

To be able to construct a reasonable BML model, exchangeability is assumed for each entity-level effect term (e.g., region and subject). Conditional on region-level effects *ξ_i_* and *ξ_j_* (i.e., when the two ROIs are fixed at indices *i* and *j*), the subject effects *π_k_* can be reasonably assumed to be exchangeable since the experiment participants are usually recruited randomly from a hypothetical population pool as representatives (hence the concept of coding them as dummy variables). As for the ROI effects *ξ_i_* and *ξ_j_*, here we simply assume their exchangeability conditional on the subject effect *π_k_* (i.e., when subject is fixed at index *k*), and address the validity of exchangeability assumption later in the Discussion.

To summarize, the main differences between the conventional GLM and BML lie in two major aspects. First, we untangle each RP-specific effect into the additive effects of the two involved regions plus interactions with a multilevel and cross-classified multi-membership structure. Second, we make the following assumption about the brain regions and RPs: the RP effects *p_ij_* under the BML models here are assigned with hierarchical Gaussian priors; in contrast, the RP effects under GLM (e.g., *p_l_* in (23)) are assumed to have a noninformative flat prior (or full independence). In other words, under the conventional GLM the effect of each RP is estimated independently from other RPs by ignoring all the interrelationships among ROIs and RPs even though the relatedness exists as captured in the correlation matrix ***P***^(*m*)^, thus there is no information shared across regions as well as across RPs. In contrast, the partitioning of each RP effect into multiple components makes it possible that the effects across regions are shared, regularized and partially pooled through a Gaussian prior under BML. It is worth emphasizing that such a cross-region Gaussian assumption shares the same rationale as, the cross-subject Gaussian assumption under the conventional ANOVA and LME.

#### MBA implementations

With the layout of crossed random effects under the LME frameworks such as LME0 (4), LME1 (12), LME2 (13), and LME3 (14), we would have to duplicate the input data and utilize both the lower (*i* > *j*) and upper (*i* < *j*) triangular parts of the IRC matrix 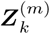 as input so that a balanced data structure (*i* ≠ *j*) can be maintained for the practical reason of typical numerical implementations (e.g., function *lmer* in *R* package *lme4* (Bates et al., 2015)). In doing so, the uncertainty estimates (e.g., standard deviations) for the obtained estimates need to be adjusted for artificially doubling the data. In addition, random-effects pairs such as *ξ_i_* and *ξ_j_* in the four LME models above as well as *ζ_ik_* and *ζ_jk_* in LME2 and LME3, are treated as two separate random-effects factors in real implementations.

For the BML systems such as BML0 to BML3 and their counterparts with covariates, we borrow the terminology and implementation strategy from multi-membership modeling. Specifically, we consider both region-specific effects of *ξ_i_* and *ξ_j_* as two members (samples or substantiations) from the same set of parameters (or the same list of regions in the neuroimaging context) with equal weights of 1, reducing the number of associated parameters from 2*m* in the LME counterpart to *m* and maintaining the original index constraint *i* > *j* in the BML models here.

#### Numerical implementations of BML

As no analytical solution is available for BML models in general, we use numerical approaches whereby we draw samples from the posterior distributions via MCMC simulations. Specifically, we adopt the algorithms implemented in Stan, a publicly available probabilistic programming language and a math library in C++ on which the language depends (Stan Development Team, 2017). In Stan, the main engine for Bayesian inferences is adaptive Hamiltonian Monte Carlo (HMC) under the category of gradient-based Markov chain Monte Carlo (MCMC) algorithms (Betancourt, 2018). The present implementations are executed with the *R* package *brms* in which multi-membership modeling is directly available (Burkner, 2017; Bürkner, 2018).

For typical BML models, examples of each of the priors (e.g., hierarchical Gaussian distributions) for cross-region and cross-subject effects as well as their interactions have been laid out in the previous section. For example, we adopt an improper flat (noninformative uniform) distribution over the real domain or a weakly informative distribution such as Cauchy or Gaussian for population parameters (e.g., *b*_0_ and *b*_i_ in (24)), depending on the minimal requirement to cope with the amount of data present; in other words, one may adopt a noninformative prior if a large amount of information is available in the data at the population level. As for assigning hyperpriors, we follow the general recommendations in Stan (Stan Development Team, 2017). Specifically, for the scaling parameters at the region and subject level, the standard deviations for the cross-region and cross-subject effects, *ξ_i_*, *ξ_j_*, and *π_i_*. as well as their interactions, we adopt a weakly informative prior such as a Student’s half-*t*(3,0,1)^4^ or half-Gaussian 𝒩_+_(0,1) (restricting to the positive values of the respective distribution). For covariance structure (e.g., **λ** in (24)), the LKJ correlation prior^5^ is used with the shape parameter taking the value of 1 (i.e., jointly uniform over all correlation matrices of the respective dimension) (Gelman et al., 2017). Lastly, the standard deviation *σ* for the residuals is assigned using a half Cauchy prior with a scale parameter depending on the standard deviation of *z_ijk_*. To summarize, besides the Bayesian framework under which hyperpriors provide a computational convenience through numerical regularization, the major difference between BML and its univariate GLM counterpart is the application of the Gaussian prior in the BML models that play the pivotal role of pooling and sharing the information among the brain regions. It is this partial pooling that effectively takes advantage of the effect similarities among the ROIs and achieves higher modeling efficiency.

Bayesian inferences are usually expressed in terms of the whole posterior distribution of each effect of interest. For practical considerations in results reporting, point estimates from these distributions such as mean and median are typically used to show the effect centrality, while quantile-based (e.g., 90%, 95%) intervals or highest posterior density intervals also provide a condensed and practically useful summary of the posterior distribution. A typical workflow to obtain the posterior distribution for an effect of interest is the following. Multiple (e.g., 4) Markov chains are usually run in parallel with each of them going through a predetermined number (e.g., 2000) of iterations, half of which are thrown away as warm-up (or “burn-in”) iterations while the rest are used as random draws from which posterior distributions are derived. To gauge the consistency of an ensemble of Markov chains, the split *Ȓ* statistic (Gelman et al., 2014) is provided as a potential scale reduction factor on split chains and as a diagnostic parameter to assist the analyst in assessing the quality of the chains. Ideally, fully converged chains correspond to *Ȓ* = 1.0, but in practice *Ȓ* < 1.1 is considered acceptable. Another useful statistic, effective sample size (ESS), measures the number of independent draws from the posterior distribution that would be expected to produce the same amount of information of the posterior distribution as is calculated from the dependent draws obtained by the MCMC algorithm. As the sampling draws are not always independent of each other, especially when MCMC chains mix slowly, one should make sure that the ESS is large enough (e.g., 200) so that a reasonable accuracy can be achieved to derive the quantile intervals for the posterior distribution. With the number of cores equal to or large than the number of MCMC chains, the typical BML analysis can be effectively conducted on any system with at least 4 CPUs.

#### BML applied to an IRC dataset

To demonstrate the modeling capability and performance of BML, we utilized a dataset from a previous FMRI study (Choi et al., 2012). Briefly, a cohort of 41 subjects (mean age = 21, sd = 2.4, 22 females) was investigated. In each of six functional runs, 169 EPI volumes were acquired with a TR of 2500 ms and TE of 25 ms. Each volume consisted of 44 oblique slices with a thickness of 3 mm and an in-plane resolution of 3 × 3 mm^2^ (192 mm field of view). The 41 subjects performed a response-conflict task (similar to the Stroop task) under safe and threat conditions (for details, see Choi et al., 2012). During all trials, after an initial 0.5-second cue signifying the beginning of each trial, there was an anticipation period during which participants viewed a fixation cross lasting 1.75 to 5.75 seconds (with duration randomly selected), after which they performed the response conflict task. Trials were separated from each other by a blank screen lasting 1.75 to 5.75 seconds (again, duration was randomly selected). During threat trials, participants received a mild shock during the anticipation period in a subset of the trials; during safe trials, shocks were never administered. Shock trials were discarded from the analysis here. To keep the trial types balanced after exclusion of physical-shock trials, the subsequent trial type after the physical-shock trial was always of the safe condition, which was also discarded from the analysis. A total of 54 trials were available for each condition. Finally, here we investigated the same 16 ROIs (listed in the first column of Table 4), as identified in the original paper (Choi et al., 2012; see also Kinnison et al., 2012).

The IRC data of 16 × 16 matrix from the *n* = 41 subjects were assembled from *m* =16 ROIs that were analyzed and discussed in Kinnison et al. (2012). With the Fisher-transformed *z*-scores of the IRC data as input, six models were performed: one GLM as formulated in (23), four BML models (BML0 to BML3, (7), (19), (20), and (21)), plus LME3 (14) that shares the same effect decompositions as BML3 (21). The GLM and BML models were fit through the *R* package *brms* (Bürkner, 2018) which is based on Stan (Stan Development Team, 2017), while the LME3 model was analyzed with the *R* package *lme4* (Bates et al., 2015). Runtime for GLM and each BML with 4 chains and 1,000 iterations (half of which were for warmup), was a few minutes on a Linux system (Fedora 25) with AMD Opteron 6376 at 1.4 GHz. For comparison, the Matlab package NBS (Zalesky et al., 2010) was used to handle multiple testing involved in the GLM approach based on the connectedness among neighboring RPs through 5000 permutations, similar to the concept of leveraging between the spatial extent (e.g., cluster size) and the effect magnitude with the overall FPR controlled at the significance level of 0.05 under NHST for voxel-wise whole brain analysis

Among the five models (GLM and four BML models), GLM yielded the worst results in terms of predictive accuracy assessed through LOO-CV (Table 2). This is most likely due to the substantial advantage BML has of characterizing the intertwined relationships among RPs instead of treating the individual RPs as independent entities under the conventional approach of massively univariate GLMs. The differences in terms of PPCs between the GLM approach and the BML models are much subtler (Fig. 3), not as clearcut as LOO-CV: the GLM tends to have generated not enough near-zero values compared to the original data than its BML competitors (cf. the peak region among the plots in Fig. 3); in addition, GLM also shows slightly worse predictions at both tails (especially the left side) beyond the inflection points.

**Table 2:** Model comparisons among five candidate models via approximate LOO-CV. Smaller values indicate better fit. To directly compare with the four BML models, the Bayesianized version of GLM (23) was fitted with the data at each RP separately. Each diagonal element (with background in blue) displays the out-of-sample deviance measured by the leave-one-out information criterion (LOOIC) and the corresponding standard error (shown within parentheses). Each offdiagonal element are the LOOIC difference between the two models (row vs. column) and its standard error (shown within parentheses). The higher predictive accuracy of the four BML models is seen here with their substantially lower LOOIC than that of GLM. Among the four BML models, two of them, BML2 and BML3, are substantially superior to the other two. Between the two most inclusive models, BML3 (with both subject-ROI and between-region interactions) is slightly better than BML2 (with between-region interaction only).

| Model | GLM | BML0 | BML1 | BML2 | BML3 |
| --- | --- | --- | --- | --- | --- |
| GLM | -2808.31 (101.65) | 1735.46 (78.20) | 3458.45 (106.53) | 1733.56 (78.03) | 3465.96 (105.94) |
| BML0 | -1735.46 (78.20) | -4543.77 (102.97) | 1722.99 (84.14) | -1.90 (0.97) | 1730.49 (84.13) |
| BML1 | -1733.56 (78.03) | 1.90 (0.97) | -4541.87 (103.04) | 1724.89 (84.21) | 1732.39 (84.15) |
| BML2 | -3458.45 (106.53) | -1722.99 (84.14) | -1724.89 (84.21) | -6266.76 (108.23) | 7.50 (7.39) |
| BML3 | -3465.96 (105.94) | -1730.49 (84.13) | -7.50 (7.39) | -1732.39 (84.15) | -6274.26 (108.22) |

**Figure 3:**
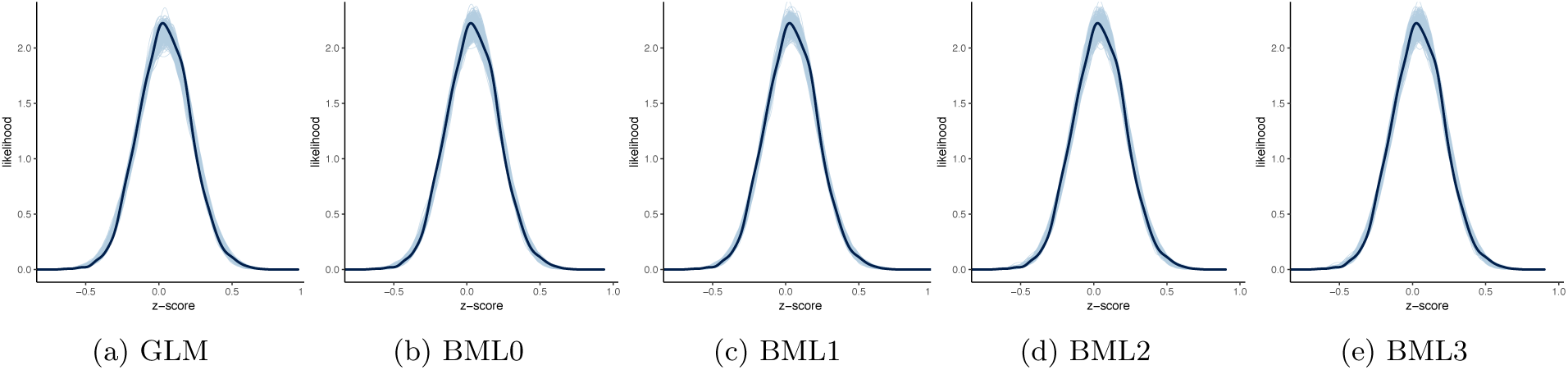
Model performance comparisons through posterior predictive checks (PPCs) and cross validations between conventional univariate GLM (*a*) and the four BML models (*b*-*e*). Each of the five panels shows the posterior predictive density overlaid with the raw data from the half off-diagonal element in a 16 × 16 Fisher-transformed IRC matrix from each of 41 subjects for the given model: solid black curve is the raw data with linear interpolation while the light blue cloud is composed of 500 sub-curves each of which corresponds to one draw from the posterior distribution based on the respective model. The differences between the solid black curve and the light blue cloud indicate how well the respective model fits the raw data. The four BML models fitted the data slightly better than the GLM (23) at the peak and both tails. However, PPC does not differentiate the five models as clearly as LOO-CV (Table 2): the visual differences in posterior predictive density with this particular dataset are small between GLM and BML and even negligible among the four BML models.

Among the four BML models, both of the most complex model BML3 in (21) and BML2 (with the subject-ROI interaction) in (20) show substantially better predictive accuracy in terms of LOO-CV (Table 2), even though all the four BML models were virtually indiscernible in terms of PPC (Fig. 3). Despite the fact that the differences between BML2 and BML3 are small as evidenced by their small LOO-CV difference and comparable standard error, we chose BML3 here for modeling demonstration and results reporting for its slight edge over BML2.

The model summary for BML3 is shown in Table 3. One noteworthy aspect is that, among the five sources of data variability, the variance for the residuals was the highest (standard deviation of 0.120 with a narrow 95% quantile interval of [0.118, 0.123]), indicating that a large amount of data variability was unaccounted for. The second and third largest sources of data variability were cross-subject effects and subject-region interaction effects, with an estimated standard deviation of 0.093 (with a 95% quantile interval of [0.072, 0.121]) and 0.070 (with a 95% quantile interval of [0.066, 0.075]), respectively; in other words, the overall variability at the subject level as well as the variability at each ROI of individual subjects are relatively large. Lastly, the variability across the 16 regions as well as the one across all RPs were relatively small with an estimated standard deviation of 0.014 (with a 95% quantile interval of [0.004, 0.026]) and 0.011 (with a 95% quantile interval of [0.004, 0.017]). Per the formulas in (17), the two ICC values for LME3 and BML3 indicate that the correlation between any two RPs of a subject that share a region was substantial with *ρ_r_* = 0.483 at the population level, while the correlation of any RP between two subjects was negligible with *ρ_s_* = 0.017.

**Table 3:** Summary results with IRC data of 16 × 16 matrix from *n* = 41 subjects fitted with LME3 and BML3 in (21). The column headers SD, QI, and ESS are short for standard deviation, quantile interval, effect sample size, respectively.LME3 shares the same effect components as BML3, and shows virtually the same effect estimate for the population mean *b*_0_ and the standard deviations for those effect components as its BML counterpart, even though in practice the input data for LME had to be duplicated to maintain the balance between the cross-random effect components. However, the LME model cannot offer statistical inferences about region- or RP-specific effects, while BML does. All Ȓ values under BML were less than 1.1, indicating that all the 4 chains converged well. The effective sample sizes (ESSs) for the population- and entity-level effects were large enough to warrant quantile accuracy in summarizing the posterior distributions for the effects of interest such as region and RP effects.

| Term | LME |  | BML |  |  |  |  |
| --- | --- | --- | --- | --- | --- | --- | --- |
| | Estimate | SD | Estimate | SD | 95% QI | ESS | $\hat{R}$ |
| $\nu$: region-subject | 0.070 | - | 0.070 | 0.002 | [0.066, 0.075] | 734 | 1.001 |
| $\lambda$: region | 0.013 | - | 0.014 | 0.005 | [0.004, 0.026] | 321 | 1.007 |
| $\tau$: subject | 0.093 | - | 0.093 | 0.012 | [0.072, 0.121] | 680 | 1.002 |
| $\mu$: region pair | 0.012 | - | 0.011 | 0.003 | [0.004, 0.017] | 366 | 1.004 |
| $b_0$: overall average | 0.041 | 0.016 | 0.040 | 0.017 | [0.005, 0.074] | 608 | 1.004 |
| $\sigma$: residual | 0.120 | - | 0.120 | 0.001 | [0.118, 0.123] | 2000 | 0.998 |

Now we start with the statistical inferences under BML3 about the 16 ROIs. As the number of regions is relatively small (*m* = 16 in this case), we can directly show the posterior distributions as well as their 50%, 90% and 95% quantiles (Fig. 4). Because the effect of “threat” condition is known from previous studies to be higher than “safe” condition, the directionality of RP effect is a priori known to be positive, and thus we focus on one-sided (e.g., positive) inferences; in other words, the quantile of a two-sided, for example, 90% quantile interval is equivalent to that of positively-sided 95% (cf. the posterior probability of the effect being positive^6^, *P*_+_, in the last column of Table 4). The posterior densities are roughly symmetric, but there are some irregularities in terms of distribution shape, especially around the peak area at regions such as BNST_L, BNST_R, Thal_R, aIns_R and SMA_R. Alternatively, the inferences on region effects can be condensed and summarized by their mean, median, standard deviation, quantile intervals, and posterior probability of the effect being positive as illustrated in Table 4. Among the 16 regions, two of them, BNST_L and Thal_R (highlighted with a green dot-dashed box in Fig. 4 and labeled green in Table 4), showed substantial region effect with strong statistical evidence as judged by the two-sided 95% quantile interval; two regions, BNST_R and Thal_L (highlighted with an orange dot-dashed box in Fig. 4 and labeled orange in Table 4), had moderate statistical evidence under the two-sided 90% quantile interval; and four regions, BF_L, BF_R, aIns_L, and aIns_R, are contralateral pairs of homotopic regions with slightly weaker statistical evidence but still sizable. These regions with sizable statistical evidence of effect were all bilateral: the basal forebrain, bed nucleus of the stria terminalis (BNST), thalamus, basal forebrain, and anterior insula, all of which are strongly involved in threat processing and wakefulness (Pessoa, 2013; Grupe and Nitschke, 2013). The BNST, in particular, is considered to be an important region during the processing of temporally uncertain threat, and has received increased attention in the past decade (Fox et al., 2015).

**Table 4:** Region effect estimates and their uncertainties under BML. Region effects at the 16 ROIs, their standard errors, 90% and 95% two-sided quantile intervals as well as the posterior probabilities of the effects being positive (the area under the posterior density with the effect being positive), *P*+, were estimated through BML3 in (21) with the experiment dataset. The lower and upper limits of the 95% (or 90%) quantile interval are listed under the columns 2.5% (or 5%) and 97.5% (or 95%), respectively. The 50% column is the median of the posterior samples. Rows in green indicate that the corresponding effect lies beyond the 95% quantile interval under BML, revealing strong statistical evidence for the region effect; rows in orange indicate that the corresponding effect lies beyond the 90% quantile interval under BML (or the 95% quantile interval if the effect sign is *a priori* known, which might be reasonable in the current context), revealing moderate statistical evidence for the region effect. Alternatively, the posterior probability of an effect being positive, *P*_+_, in the last column can be used as statistical evidence. For example, *P*_+_ = 0.942 at Thal_L indicates that there is some extent of statistical evidence for the region’s effect as the posterior probability of the effect being negative is 0.058. In fact, the statistical evidence for the four regions of BF_L, BF_R, aIns_L, aIns_R and SMA_R are quite substantial based on their posterior probabilities *P*+. The conventional GLM would not be able to make direct inferences about individual regions for MBA. Unlike the popular practice of sharp thresholding under NHST, more customized quantile intervals (e.g., 10%, 50% and 90%), if desirable, can be added in the final reporting in order to make corresponding inferences. More importantly, the effect magnitude and the associated uncertainty provide richer information and more accurate quantification for the relative importance of a region than the concept of “hub” under the neuroimaging graph theoretic analysis through counting the number of “surviving paths” associated with a region, using some extent of binarization for the RPs. For example, the difference between the column of “mean” and that of 50% quantile (median) is an indicator of skewness. Region abbreviations are as follows. BF: basal forebrain; BNST: bed nucleus of the stria terminalis; IFG: inferior frontal gyrus; IPG: inferior parietal gyrus; Ins: insula; MPFC: medial prefrontal cortex; SMA: Supplementary motor area; Thal: thalamus. Other abbreviations: a: anterior; p: posterior; m: medial; L: left; R: right.

| ROI | mean | std err | 2.5% | 5% | 50% | 95% | 97.5% | $P_+$ |
| --- | --- | --- | --- | --- | --- | --- | --- | --- |
| BF_L | 0.026 | 0.017 | -0.008 | -0.001 | 0.026 | 0.055 | 0.060 | 0.942 |
| BF_R | 0.024 | 0.016 | -0.007 | -0.003 | 0.024 | 0.051 | 0.057 | 0.924 |
| BNST_L | 0.032 | 0.017 | 0.001 | 0.005 | 0.032 | 0.061 | 0.067 | 0.977 |
| BNST_R | 0.029 | 0.017 | -0.002 | 0.002 | 0.028 | 0.057 | 0.062 | 0.960 |
| Thal_L | 0.030 | 0.017 | -0.004 | 0.001 | 0.029 | 0.059 | 0.064 | 0.952 |
| Thal_R | 0.036 | 0.019 | 0.000 | 0.006 | 0.035 | 0.067 | 0.073 | 0.976 |
| aIns_L | 0.025 | 0.017 | -0.006 | -0.001 | 0.025 | 0.053 | 0.060 | 0.944 |
| aIns_R | 0.025 | 0.017 | -0.008 | -0.003 | 0.025 | 0.054 | 0.059 | 0.937 |
| IPG_L | 0.011 | 0.017 | -0.024 | -0.018 | 0.012 | 0.039 | 0.045 | 0.758 |
| IPG_R | 0.014 | 0.017 | -0.021 | -0.015 | 0.014 | 0.041 | 0.047 | 0.794 |
| MPFC_L | 0.008 | 0.017 | -0.030 | -0.021 | 0.009 | 0.036 | 0.040 | 0.694 |
| MPFC_R | 0.010 | 0.017 | -0.026 | -0.019 | 0.011 | 0.038 | 0.042 | 0.730 |
| mIns_R | 0.012 | 0.017 | -0.023 | -0.017 | 0.012 | 0.039 | 0.045 | 0.764 |
| pIFG_L | 0.005 | 0.019 | -0.032 | -0.026 | 0.006 | 0.035 | 0.039 | 0.622 |
| pIFG_R | 0.016 | 0.017 | -0.018 | -0.011 | 0.017 | 0.043 | 0.049 | 0.838 |
| SMA_R | 0.021 | 0.016 | -0.010 | -0.006 | 0.021 | 0.047 | 0.052 | 0.908 |

**Figure 4:**
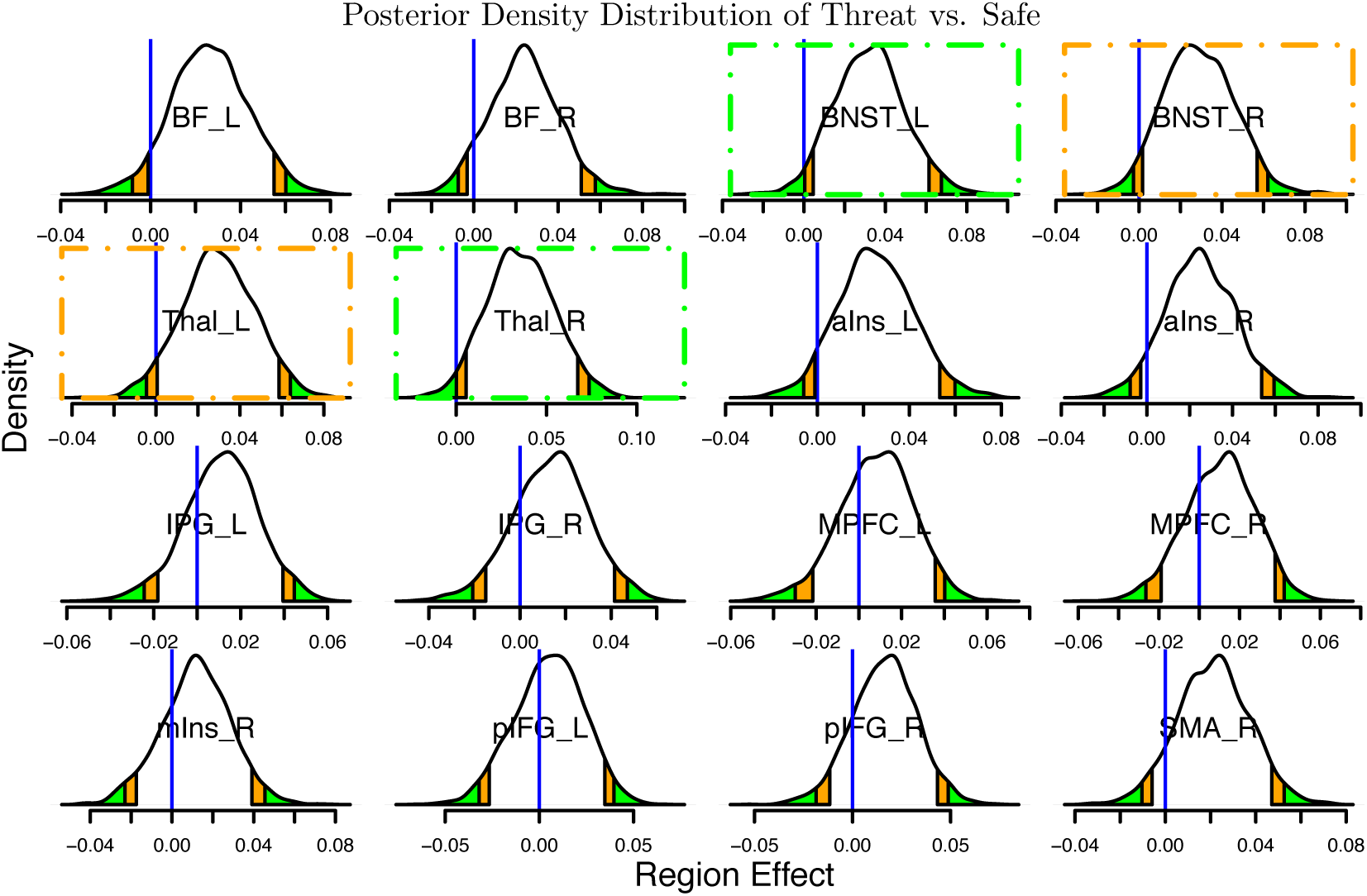
Posterior density plots of region effect for threat versus safe based on 2000 draws from BML3 in (21) with the experiment dataset. The blue vertical line indicates zero region effect, orange and green tails mark areas outside the two-sided 90% and 95% quantile intervals, respectively. The two ROIs (BNST_L and Thal_R, labeled with a green dot-dashed box) with strong statistical evidence of region effect (based on 95% quantile intervals) can be identified as the blue line being within the green tails, while two other ROIs (BNST_R and Thal_L, labeled with an orange dot-dashed box) show moderate statistical evidence of region effect (the blue vertical line lies within the orange band). Four more ROIs that form two contralateral pairs of homotopic regions, BF_L and BF_R, aIns_L and aIns_R, also contain some extent of statistical evidence as they are close to the typical threshold adopted for discussion convenience. The posterior density provides much richer information about each effect such as spread, shape and skewness; unlike the conventional confidence interval that is flat and inconvenient to interpret deeply, it is valid to state that, conditional on the data and model, we believe that with probability, for example, 95% the region effect is in its 95% posterior interval. The two-sided 90% quantile interval can be interpreted as one-sided 95% if the effect directionality is a priori known.

To recapitulate, BML offers a unique feature of statistical inference for region effect that is not available with the conventional GLM. The effectiveness of estimating region-level effects under BML is illustrated by the results obtained with the dataset investigated here (Choi et al., 2012). Specifically, BML identified the BNST and the thalamus as exhibiting region-level effects (stronger statistical evidence was obtained particularly for the left BNST and the right thalamus, but the respective contralateral sides also exhibited moderate statistical evidence). The identification of the BNST is particularly noteworthy because of its involvement in processing threat during uncertain and more temporally extended conditions. For example, in a previous threat study, the “betweenness” (threat vs. safe periods) of the BNST was modulated by anxiety scores, such that greater increases in betweenness were observed for participants with high- relative to low-anxiety (McMenamin et al., 2014). The thalamus is also a key region in the processing of threat, and is at the core of cortical-subcortical signal integration that is required for determining the biological significance of stimuli and contexts (Pessoa, 2017).

In contrast, presenting RP results is more challenging. As the number of RPs 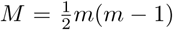 is usually quite large (e.g., 120 in the current dataset), it might not be practical to show the full results in details in the format of density histograms (Fig. 5) or in the format of summarized results with mean, standard error and quantile intervals as illustrated for the region effects in Fig. 4 and Table 4. Instead, one may present the results with a matrix format that is typically seen in the literature, that is, a more condensed gamut of RP results with the effect estimates color-coded in a symmetric matrix (e.g. side by side comparison of GLM and BML3 in Fig. 6), revealing their strengths with some extent of statistical evidence (e.g., based on quantile intervals).

**Figure 5:**
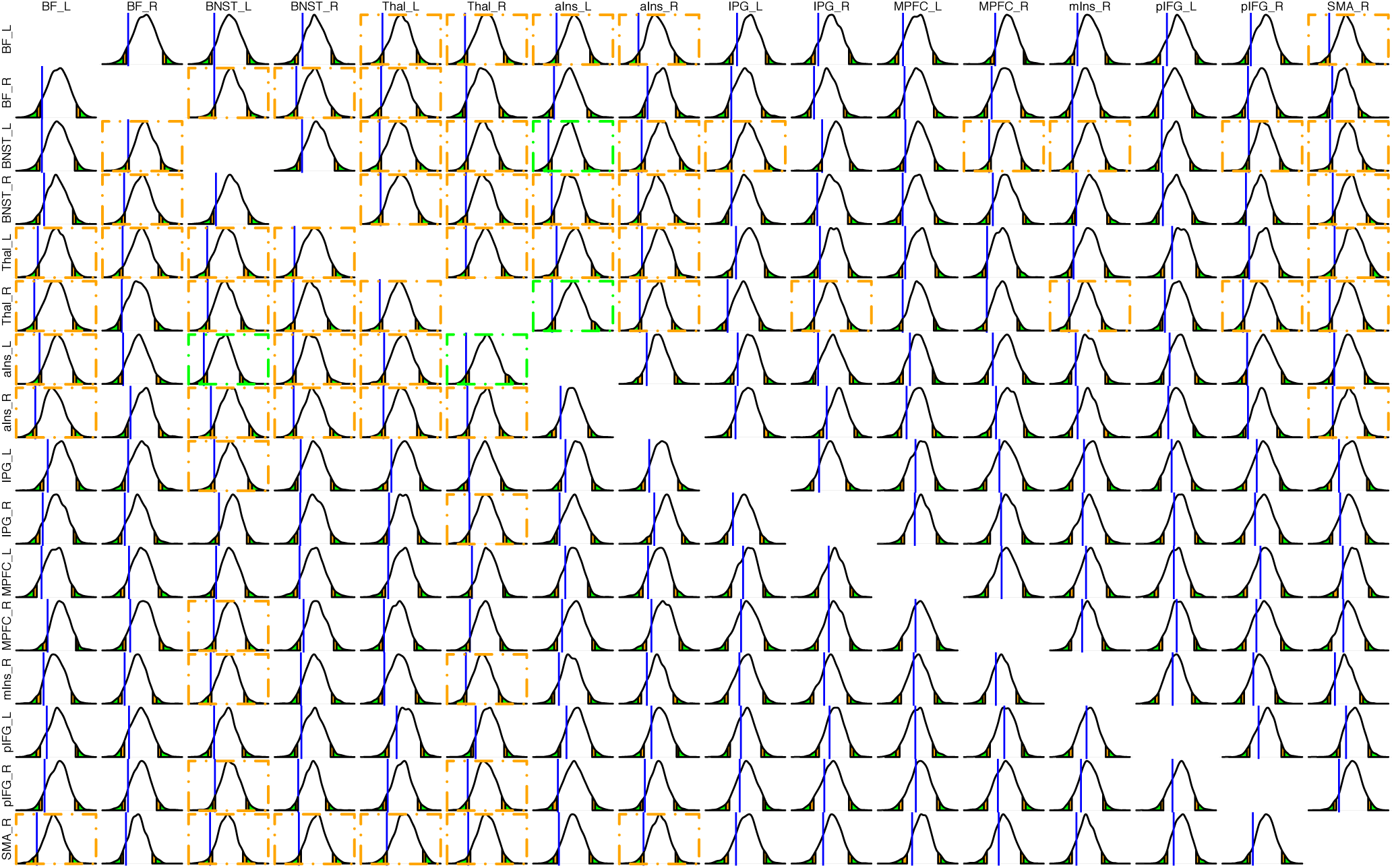
Posterior density plots for the effect magnitude of threat vs. safe at all RPs for BML3 (21). As in Fig. 4, the blue vertical line marks the location of zero effect, orange and green areas under the density curve show the ranges outside the 90% and 95% quantile intervals, respectively. RPs with some extent of statistical evidence are labeled with an orange or green dot-dashed box based on their two-sided 90% or 95% quantile intervals. The empty slots along the diagonal correspond to the correlation value of 1 along the diagonal of the IRC matrix. A condensed (and perhaps more straightforward) version of the RP effects for this model is presented in Fig. 6 (right).

The comparisons of RP effects between GLM and BML can be summarized as follows. First of all, the effect of partial pooling or shrinkage is evident relative to GLM from the following perspectives: the effects at both large and small (as well as positive and negative ends) estimated under the GLM tend to be “dragged” toward to the center under BML, as evidenced by large (darker) or small (lighter) circles in GLM (Fig. 6, left) corresponding to slightly smaller (lighter or above zero) or larger (darker) under BML (Fig. 6, right). In addition, GLM initially identified 62 RPs (Fig. 6, left), among the total 120 RPs, as statistically significant under the one-sided NHST level of 0.05. However, none of them survived the correction for multiple testing through the permutation approach implemented in the Matlab toolbox NBS (Zalesky et al., 2010). As a direct comparison, 33 RPs from BML3 (Fig. 6 right) are considered to have effect with an equivalent extent of statistical evidence (i.e., based on a comparable 95% one-sided quantile intervals), and they form a subset of those 63 RPs identified under GLM without multiple testing correction.

**Figure 6:**
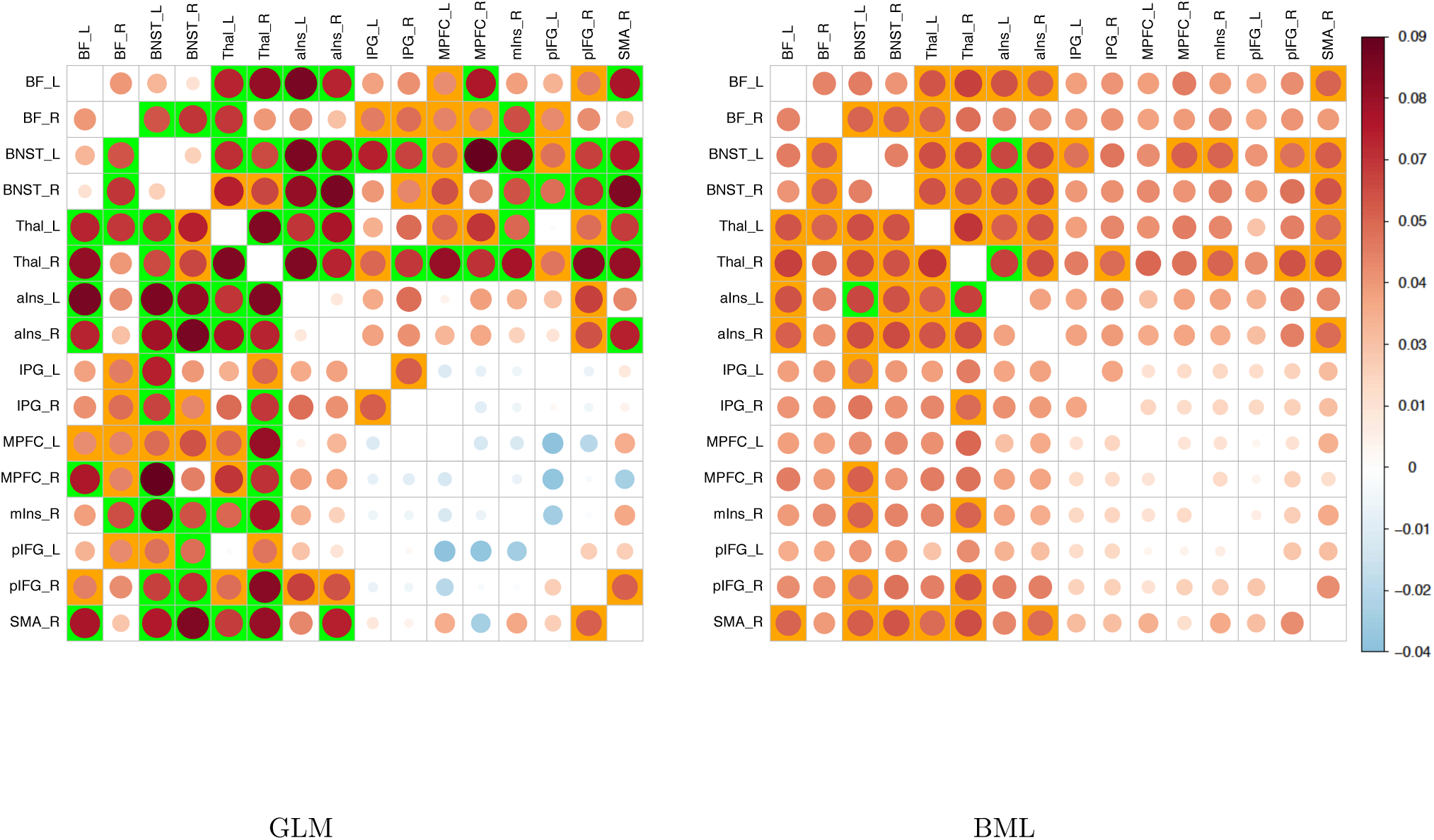
Comparisons of inferences on RP effects between GLM in (23) on the left and BML3 in (21) on the right with the experiment data. The empty slots along the diagonal are trivial cases that correspond to the correlation value of 1 along the diagonal of the IRC matrix. The effect magnitude of threat vs. safe is symbolized with both circle size and color scheme (colorbar, far right). The impact of partial pooling or shrinkage under BML is evident in the sense that the effects for most RPs are “dragged” toward to the middle relative to their counterpart under GLM. The RPs with strong statistical evidence of effect based on two-sided 95% confidence (or quantile) interval are colored in green while those with moderate evidence based on two-sided 90% confidence (or quantile) interval are labeled in orange. As the directionality of the effect is a priori known (i.e., positive for the contrast of “threat” relative to “safe”), one-sided effect is considered here, therefore two-sided 90% and 95% confidence (or quantile) intervals shown in the color coding are equivalent to one-sided 95% and 97.5% intervals, respectively. While there are 63 RPs (both green and orange) that are identified as statistically significant under the significance level of one-sided 0.05 without correction for multiple testing, 33 RPs can be considered as having effect with moderate evidence if a comparable criterion is adopted under the Bayesian framework. None of the RPs under GLM were rendered as statistically significant at the “cluster” level FPR of 0.05 through permutations in NBS under one-sided testing, while multiplicity handling is embedded in the BML model through the constraint (or partial pooling) of Gaussian priors. It is worth emphasizing the importance of full results reporting as well as the avoidance of dichotomous interpretation. The choices of highlighting the two quantiles, 90% and 95%, are purely for the convenient comparisons with the conventional thresholding under GLM, but it is more important to present the full results even if the investigator prefers to practically focus on a limited list of effects based on a particular cutoff.

The inefficiency and disadvantage of the conventional GLM, compared to BML, is well illustrated here with this experiment dataset. On one hand, a substantial number of RPs were initially labeled under GLM as “statistically significant” with the presumption of all RPs being unrelated. On the other hand, none of the RPs would be claimed to be “statistically significant” once a correction procedure, NBS, was applied to rectify the multiple testing issue due to the incorrect presumption. In contrast, BML integrates all effects into one model, quantitatively capturing the scaffold of relatedness among the various effect components. As multiplicity is directly handled through partial pooling under BML, no further “correction” for multiple testing is needed (Gelman et al., 2012), achieving a relatively higher inference efficiency.

The scientific relevance of those RPs with some extent of statistical evidence under BML is as follows. Given, as noted, the involvement of the basal forebrain, BNST, thalamus, and anterior insula in threat processing, our a priori expectation was that RP effects would increase during the threat relative to the safe condition when these regions were involved (inferentially, one-sided assessment is therefore appropriate). For multiple RPs involving the above regions, their effects exhibited substantial increases during threat, consistent with the notion that they form a more tightly cohesive cluster composed of the basal forebrain, BNST, and thalamus during this condition (Kinnison et al., 2012).

#### Discussion

Network modeling is popular and widely adopted in neuroimaging because it allows the investigator to probe interregional information flow in the brain. The popularity is evidenced by the explosively growing number of papers in literature that explore the differences between controls and a disease group (autistic, Alzheimer’s, Schizophrenic, depressive, etc.) through MBA. However, the dangers are also obviously omnipresent: model inefficiency and inconsistency, the involvement of arbitrary decision rules (e.g., thresholding), artificial discretization of IRCs, and over-interpretation may loom large when the models are applied without being rigorously checked, compared and validated.

At present, there are broadly two major approaches to handling MBA in neuroimaging. The first one adopts the typical GLM approach, similar to the whole-brain voxel-wise group analysis, and this has been implemented into various software packages such as the Matlab toolbox NBS (Zalesky et al., 2010) and FSLNets in FSL (Jenkinson et al., 2004). The second approach is the adoption of graph theory into neuroimaging through several variations (e.g., Bassett et al., 2018), which mainly focuses on topological properties derived from treating brain regions as connected entities in a network.

##### MBA: current approach one - network-based statistics (NBS) through GLM

The fundamental issue of massively univariate GLM is its inefficiency in statistical inferences. As a time-honored analytical tool, GLM is easy to understand and adopt. Its underlying mechanism bears the same hallmark of conventional whole-brain voxel-wise neuroimaging data analysis; that is, each spatial element (voxel) is treated as an independent entity from its neighbors (even though spatial correlation is present in reality). At the ROI level for IRC data, the same GLM strategy applies with the voxel replaced by the RP, but the spatial relatedness for IRC is more intricate, as shown in the correlation structure ***P***^(*m*)^ as illustrated in Fig. 2, than the relatively simple spatial relationship among neighboring voxels in whole-brain analysis (typically characterized through a single number: full width at half maximum). This fundamental assumption for the massively univariate modeling methodology provides a convenient platform to apply for conventional analytical tools such as GLM, but one severe consequence is the issue of multiplicity due to the pretense of unrelatedness during model building: the analyst has to deal with the penalty from multiple testing because the same model is applied as many times as the number of entities (voxels for whole brain or RPs in the case of MBA). The GLM approach for MBA face the same hurdle in dealing with the correction issue as the typical the whole-brain voxel-wise analysis does (Chen et al., 201a). It is this modeling inefficiency that levies a heavy penalty on inference efficiency, as demonstrated here by the performance of multiple testing correction through permutations on the GLM methodology with the experiment dataset.

As a means for correcting for multiple testing in whole-brain voxel-wise analysis, permutation testing has been applied to the context of MBA through NBS (Zalesky et al., 2010). Specifically, NBS borrows the concept from an early version of permutation testing in voxel-wise data analysis (Nichols and Holmes, 2001), in which the construction of a null distribution of a maximum statistic (either maximum testing statistic or maximum number of surviving RPs that form a cluster based on a predetermined threshold for the testing statistic) through permutations. The original data are assessed against the null distribution, and the top winners selected at a designated rate (e.g., 5%) among the testing statistic values or “connected” RPs are declared as the surviving ones (Zalesky et al., 2010).

The application of permutation testing to MBA has had several challenges to try to address, even though its counterpart is relatively effective in maintaining the nominal FPR level for voxel-wise data.

1. Arbitrary thresholding. In NBS, a primary threshold has to be preset for the testing statistic in the case of maximum cluster size of “connected” RPs. However, there is no principled way to allow the investigator to choose such an initial threshold.
2. Garden of forking paths. Arbitrary threshold leaves the door open for a trial-and-error approach^7^, resulting in a potential problem for the multiplicity issue of “forking paths,” especially when the final results sensitively depend on different primary thresholds: a slightly different primary threshold may end up with a different set of clusters or RPs with different properties. Even though each study usually reports only one of many potential results in the literature, the number of trials are most likely hidden from the publication. Nevertheless, evidence of arbitrariness in primary thresholding leading to different results can still be seen under NBS (Baggio et al., 2018). For example, with four primary thresholds at one-sided p-value of 0.001, 0.005, 0.01 and 0.05, permutation testing in NBS rendered in a recent study (Yin et al., 2018) dramatically different results: 8, 41, 78 and 188 RPs, respectively, after family-wise error correction.
3. Failure to capture the intricate relatedness among RPs. The interrelationships among regions and RPs are not directly modeled in NBS’s GLM approach; in the end, it is only the number of RPs that matters in the step of multiple testing correction through the definition of clusters among RPs. A side effect of this is that the “importance” of a region (its degree or “hubness”, in graph theoretic terms) is judged solely by the number of surviving RPs, not by the effect at each region, making it further sensitive to the specifics of primary thresholding.
4. Inference inefficiency. As a consequence of ignoring the interrelationships among regions and RPs, the penalty of GLM from the correction step can be very severe, resulting in inference inefficiency as demonstrated with the experiment dataset here.
5. Inappropriate modeling in the presence of within-subject variables. When one or more within-subject variables are involved, the error structure under univariate GLM has to be properly stratified so that each testing statistic (e.g., *F*- or Student *t*-statistic) can be correctly formulated with the corresponding denominator. However, the NBS implementations (Zalesky et al., 2010) have adopted the same univariate GLM strategy in handling within-subject variables as in some packages of whole-brain voxel-wise analysis, where the residual term is uniformly utilized for all statistic formulas regardless of the nature of the variable type (between- or within-subject). Such mishandling of within-subject variables has been shown to lead to problematic inferences for some effects (Chen et al., 2014).
6. One-sided testing. A pair of one-sided testing is adopted in NBS as default as commonly practiced in neuroimaging, leading to an undesirable and unnecessary doubling of false positive rate (Chen et al., 2018b).

To contain the arbitrariness involved in the primary permutation version implemented in NBS, a modified version of permutation testing for NBS has been recently proposed (Baggio et al., 2018). The modification, termed as threshold-free network-based statistics (TFNBS), is an extension of its whole brain analysis counterpart, threshold-free cluster enhancement (TFCE) (Smith and Nichols, 2009), that takes an integrative consideration between signal strength and spatial relatedness. The TFNBS approach hinges on two parameters that control the integration between the effect magnitude of each RP and the cluster size, and those two parameters have to be predetermined for all scenarios. In effect, the arbitrariness for primary thresholding is converted and mapped to the specification of these two empirical parameters that are associated with spatial extent and effect strength, respectively. While TFCE is relatively successful in effectively maintaining the overall FPR at the whole brain level by narrowing down the choices of the two parameters within limited ranges, the effectiveness of TFNBS remains questionable: the varying ranges of those two parameters remain relatively wide. It is unclear as to what is the main cause of performance differences between TFCE for voxel-wise data analysis and TFNBS for MBA, but one possible issue might be the difference of spatial correlation structure between whole-brain data and IRC matrix. For the whole brain data, the spatial relatedness is implicitly embedded in the contiguous supporting section defined through the TFCE formula, which is relatively effective. However for TFNBS, the spatial relations among RPs, as characterized in ***P***^(*m*)^, are fully disregarded, and are only mapped to discretized elements under NBS as a form of counting surviving RPs. In other words, all that matters in the end is the “cluster” size in terms of supra-threshold RPs. This may lead to a large number of topological variations (e.g., Fig. 7) and much wider ranges in tuning the two parameters than its conventional counterpart TFCE for whole-brain data analysis. The inefficiency of GLM for MBA combined with permutation testing is likely related to the fact that, when generating null distributions of clusters with “connected” surviving RPs, the topological differentiations are not taken into consideration. In essence, the fundamental issue with NBS and its modified version TFNBS might be that the relatedness structure among the IRC matrix elements is neither incorporated in the model nor maintained in randomly generating the null distribution through permutations.

**Figure 7:**
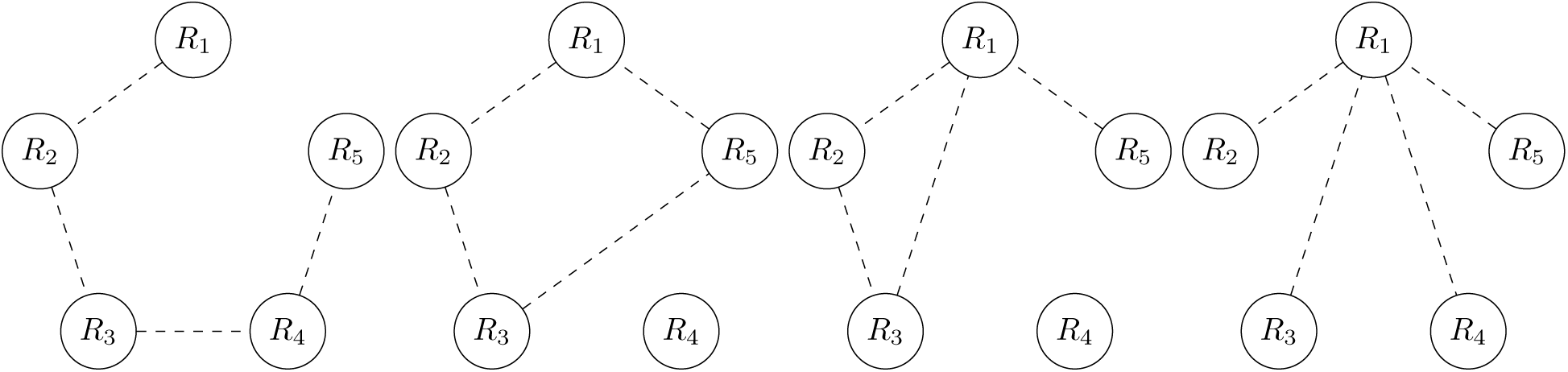
Four different IRC scenarios with four “surviving” RPs through the practice of thresholding among *m* = 5 ROIs. Unlike the typical representations with solid lines in the field, we intentionally adopt dashed lines here to indicate correlations, not real connections, between regions to avoid potential misinterpretations. Even though these four scenarios have the same number of “surviving” RPs after thresholding and thus are not differentiable from the perspective of cluster size or density, their topological structures are quite different; for example, region *R*_1_ involves 1, 2, 3, and 4 “surviving” RPs for the four scenarios, respectively. These four cases are simply for demonstration purpose because they are not an exhaustive list of possibilities for four “surviving” RPs among five ROIs, and the number of cases increases dramatically as the number of regions and that of RPs become large.

In summary, unlike in the whole brain analysis in which the spatial relationship is considered when handling multiplicity, the GLM-based permutation testing for MBA does not take into consideration the intricate relationships among the regions and among the RPs as demonstrated here. As a result, the GLM method, when dealing with multiplicity correction, tends to suffer from excessive penalties, loses inference efficiency, and depends sensitively on one or more parameters in the currently available implementations.

##### MBA: current approach two - graph theoretic analysis

A second popular approach for MBA is the fast-growing applications of graph theory, which aims to investigate synchronization, communication, coding and information transmission in a neural network. The motivation is that, at a specific level (e.g., neuron, anatomical region), information flow or neuronal signal transmission in the brain can be abstractly approximated as ensembles of networks (and subnetworks) represented with the presence and absence of connections among involved regions. Therefore, the IRC matrix, as one of the few data extractions, could be utilized to extract various topological properties among brain regions.

While graph theory methods contain sophisticated mathematics and terminology, most of the graph theoretic analyses in neuroimaging are largely built on a binarized approach through substantial thresholding and dichotomization. For example, a typical strategy is 1) to determine whether a RP is included in the surviving network through the threshold value of, for example, 0.2 for the correlation value, or 2) to limit the number of surviving RPs based on a density cutoff of, for example, 15%. Thereupon a long list of graph measures are explored through counting the number of surviving RPs and differentiating the fine-grained topological structures of the surviving graph (Medaglia, 2018). For instance, at the micro-scale level, the concept of degree for a region in a surviving graph is defined as the number of surviving RPs associated with a region, leading to other measures such as hub coefficient, node centrality, node efficiency, betweenness centrality, closeness centrality, cliques, holes or cavity, clustering coefficient (or local efficiency), eigenvector centrality, paths, and shortcuts. At the mesoscale level, the surviving graph is divided into multiple modules or communities, resulting in terms such as modular structure or community organization, rich club, network core, core-periphery structure or assortative organization. Lastly, at the macro-scale level, single values are typically used as measures representing the properties of a surviving graph, and such global measures include graph density, preferential attachment, region fitness, global clustering coefficient, characteristic path length, global efficiency, small-worldness, graph communicability and minimum spanning tree. In short, there is a large list of topological parameters to explore and interpret.

It is beyond the scope of the current context to evaluate the full merits and disadvantages of this widely adopted methodology. Here we simply raise a few concerns from our perspective.

1. Arbitrary thresholding. A substantial amount of graph theoretic work relies on strict and sharp thresholding and/or binarization. The fundamental question is, how rigorously do human brains follow such a preset value based on convenience (threshold of, say, 0.2, but not 0.187)? To date, there does not appear to be an established rigorous threshold that is associated with physiological reality. Even under NHST, the rejection of a null hypothesis (e.g., failure to pass the correlation threshold of 0.2 or RP density of 15%) for a particular RP does not necessarily mean the nonexistence of the RP effect. By the same token, if the failure of an RP to survive a threshold does not necessarily mean the nonexistence of the effect, how useful are a myriad of concepts such as holes or cavities that are defined as “the absence of connections?”
2. Disregard of uncertainty. It remains unclear how much sensitivity exists for each step when the involved threshold changes in the process of extracting those condensed topological metrics: no uncertainty information (e.g., standard error for the number of surviving RPs) is ever explored even though statistical quantities are utilized. When a different threshold is adopted, is it possible that a rich club would be no longer rich, and a small world might not be small any more? Statistics is the science of dealing with uncertainty, and that uncertainty is central to interpreting and weighing the final results. Without uncertainty available as part of the final results, any characterization such as the long list of metrics from graph theoretic analysis can be easily misinterpreted as deterministic.
3. Garden of forking paths. When a cascade of multiple steps is involved in deriving a graph measure, how many forking paths and ramifications may be possible, particularly when there are so many topological metrics to choose from (e.g., hub coefficient, node centrality, rich club, network core, global efficiency and dozens more)? The number of steps, substantially magnified by the large number of metrics, exacerbates the severity of “forking path,” and reduces the reliability and robustness of the final results and interpretations. Furthermore, no correction has been formally adopted in the field to handle the multiplicity issue involved with the topological metrics that are simultaneously computed in each study.
4. Oversimplification through data extraction. Dimensional reduction is a useful information extraction. Nevertheless, it is important to recognize the limitations and information loss of each extraction due to oversimplification. For example, the global parameters (characteristic path length, global efficiency, modularity, small-worldness, and assortativity coefficient) in graph theory tend to suffer from specificity during data reduction. As the problems with arbitrary threshold are recognized, there have been some efforts to modify or improve the information extraction process. For example, a “weighted,” instead of binarized, approach is occasionally adopted at the RP level of each IRC matrix in graph theoretic analysis. However, artificial data manipulations are still involved through hiding the negative IRC values - changing them to zero or their absolute values. In addition, the uncertainty information (e.g., standard error) after averaging across subjects is typically discarded in the later steps. Another method modification is to adopt multiple thresholds instead of a single one so that a wide range of explorations could be conducted to check the sensitivity of those graph metrics. However, a resulting issue is another potential route leading to forking paths and multiplicity. Since even the mapping of IRC measures to physiology is uncertain, the mapping of the popular terms and topological notations such as edge, path, shortcut, core-periphery, etc. could also be problematic, particularly when their values are so threshold-dependent.

##### General comments about BML in handling MBA

For communication convenience, we note that our usage of region or ROI corresponds to node or vortex in network modeling and graph theoretic analysis; RP to path, edge, or connection; and the relative strength of RP effect to hubness. The adoption of distinct terms is mainly due to the consideration that relationships across regions and across RPs are characterized through effect decompositions under the BML framework. The focus thus is on the quantitative characterization of various effect components such as region- and RP-specific effect estimations and their uncertainties. In addition, even though the investigator may choose a particular quantile interval under BML as a threshold for highlighting strong evidence of the effect of interest, dichotomization based on a cutoff value is not likely an accurate characterization for the inter-region relationships. Therefore, we strongly suggest full results reporting instead of building up a network through thresholding. Lastly, even if we ignore the subtlety of Pearson correlation coefficients measuring signal similarity, broader modeling exists so that it still might not be an accurate mapping to assign all effects being studied (e.g., the association between age and RP effects) to the notion of edges, paths, or connections.

Through the decomposition of each matrix element into additive effects of multiple components under BML, we can more accurately capture the intricate relationships among regions as well as among RPs just using the data itself, and several effects of interest can be directly derived through posterior distributions without having to resort to hard thresholding decision. In addition, the uncertainty contained in the posterior distribution of an effect naturally propagates to all quantities computed based on the posterior distribution. For example, the effect and its uncertainty at each region and each RP can be conveniently retrieved as illustrated in Table 4, Figures 4, 5 and 6. In addition, the relatedness between any two RPs that are associated with a common region can be quantified through an ICC value, as demonstrated by *ρ_r_* = 0.483 (cf., *ρ* in Fig. 2) with our experiment dataset.

We believe that the typical RP-wise GLM approach for MBA is inaccurate and inefficient because the common information shared across regions and RPs are fully ignored even though a final “network” is intended to be inferred from the analysis. Instead, we propose a more efficient approach through BML that could be used to confirm, complement or even replace the GLM methodology. To derive the approach, we started with a population analysis strategy with ANOVA or linear mixed-effects (LME) by incorporating each RP as two crossed random-effects components relative to each of the two involved regions in addition to the subject-specific component. Then we translated the LME model to a Bayesian framework, and used this BML framework to resolve the issue of multiple testing involved in the RP-wise GLM approach. The proposed BML approach dissolves multiple testing through more accurately accounting for both data structure and the shared information across regions and RPs, and consequentially improving inference efficiency. In a nutshell, the crucial feature here is that the RPs are not assumed to be unrelated as under the conventional GLM, but instead are treated as being associated with each other through a Gaussian distribution assumption under BML.

One aspect of Bayesian modeling we should comment on is the notion of “subjectivity” of priors that is often brought up by statistics consumers who were largely trained under the conventional NHST framework. It is beyond the scope of the current paper to discuss all details and advances within the Bayesian field (see, for example, Gelman and Hennig, 2016). However, we would like to point out that the major prior for the cross-region components, *ξ_i_* and *ξ_j_* in the BML models here, for example, is just a Gaussian distribution, which is not so different from the same type of assumption placed on the cross-subject variability and residuals under the conventional statistical model such as regression, AN(C)OVA, GLM, and LME. The impact of priors for other parameters (e.g., intercept or population effect in the BML model and the variances for the priors) is usually negligible if the amount of data is reasonably large. In short, the priors and hyperpriors typically adopted in BML are weakly informative and have very little influence on final results. On the other hand, with hyperpriors, BML can solve a system that would be considered over-parameterized and over-fitted under, for example, GLM or LME; more importantly, it can make statistical inferences that would not be feasible under the conventional framework. For researchers who are worried about using informative priors, it is always possible to specify very wide non-informative or weakly informative priors.

Some subtle BML features also warrant detailed discussion here. Even though the investigator may have to focus on a list of results (e.g., regions or RPs) based on a preset criterion, the existence of such a choice for convenience (e.g., highlighting certain regions or RPs above some threshold) does not indicate a real, underlying black-and-white distinction between those surviving effects and the ones that fail to pass the filtering funnel, nor does it mean the existence of such distinction in reality (e.g., similarity of BOLD response patterns between two regions). That is, through thresholding, the investigator may highlight a limited set of results with relatively strong statistical evidence, but the effects themselves do not depend on the particular threshold. Therefore, the investigator can still present all the unfiltered results, e.g., in an Appendix or supplementary material. For example, the effects at two contralateral pairs of homotopic regions, BF_L and BF_R, aIns_L and aIns_R, showed some statistical evidence that is close to the conventional threshold (Fig. 4). Similarly, the effects for RPs such as BNST_L and BNST_R, and BF_L and BF_R exhibited some extent of statistical evidence (Fig. 5), as they were rather close to the (arbitrary) threshold cutoff adopted for discussion convenience here. From a neurobiological standpoint, this is quite reasonable given that these pairs involve corresponding regions across the hemispheres, which often exhibit correlated responses. Similarly, the effects of RPs such as Thal_R and BF_R, and BNST_L and BF_L also exhibited statistical evidence (Fig. 5) that was very close to the threshold; this statistical evidence is consistent with the fact that correlated responses in the same hemisphere is often observed for regions that are functionally related in particular contexts (here, threat). These observations leave the door open for further substantiation in future studies instead of being marked as “non-existent” in a dichotomous framework.

One may notice that some negative effect estimates under GLM (particularly with a small magnitude) correspond to positive values under BML. Such negative effects would typically be zeroed or considered nonexistent under graph theoretic analysis. However, we emphasize that Bayesian inferences do not rely on point estimates to the same degree that conventional statistics do; that is, the point estimate for an effect should not be isolated from its uncertainty or quantile interval under the Bayesian framework. For example, using the three RPs between pIFG_L and each of the three regions MFPC_L, MFPC_R and mIns_R, as an example, even though their point estimates were negative under GLM (−0.037, −0.036 and −0.031, respectively, Fig. 6 left) and positive under BML (0.005, 0.006 and 0.008, respectively, Fig. 6 right), their 95% quantile intervals under BML are [-0.061, 0.071], [−0.060, 0.072] and [−0.058, 0.074], respectively, indicating that these three effects could be negative with almost equal likelihood.

For BML modeling with no between-subject explanatory variables, the most inclusive model BML3 tends to perform well, as demonstrated with the experiment dataset here. However, when at least one between-subject explanatory variable is present, we recommend a BML model such as (24) without the interaction effects explicitly modeled between regions and subjects. Specifically, if the interaction effects, such as *ζ_ik_* and *ζ_jk_* in BML2 in (20) and BML3 in (21), were included in the BML model (24), cross-subject variability at each region would be largely explained by the interaction effects *ζ_ik_* and *ζ_jk_*, leaving little for the effect of interest, *ξ*_1*i*_ and *ξ*_0*j*_. Therefore, when the region-specific effect associated with a between-subject explanatory variable (e.g., group difference or age effect at each ROI) is the research focus, it is a prerequisite that such effect be not substantially absored by the interaction effects between regions and subjects.

##### Advantages of BML in handling MBA

Our adoption of BML for MBA, as illustrated with both theory and real experimental data, indicates that BML offers the following advantages over traditional GLM and graph theoretic analysis:

1. Generality. As BML and LME usually share a corresponding modeling structure, BML can handle data structures involved in the conventional models that are subsumed under LME such as Student’s t-tests, ANOVA, regression, ANCOVA and GLM. For example, missing data can be handled as long as the missingness can be considered random; therefore BML can be reasonably applied to SAM analysis with DTI data in consideration of some fraction of a group not having some white matter connections found, for example, due to noise. BML is superior to LME in dealing with complicated data structures. For example, the number of parameters under LME with a sophisticated variance-variance structure could be high, leading to overfitting and convergence failure with the maximum likelihood algorithm; in contrast, the priors under BML may help overcome the overfitting and convergence issues.
2. Hierarchization. The crucial feature of BML (and its LME counterpart), when applied to MBA, is the disentangling of each RP effect into the additive effects of the two involved regions plus other interaction effects. Thanks to this untangling process, both region- and RP-specific effects can be retrieved through their posterior distributions under BML, which would not be achievable under LME. The reason is that, as each effect under LME is categorized as either fixed- or random-effects component, only the fixed-effects components (e.g., *b*_0_ in LME0, LME1, LME2, and LME3) can be inferred with effect estimates and uncertainties while random-effects components (e.g., region effect *ξ_i_*, RP effect η_*ij*_, interaction between region and subject *ζ_ik_* in LME0, LME1, LME2, and LME3) would only be assessed with their variances. In contrast, as all effect components are considered random under the Bayesian framework, they all can be, either directly or through reassembling, estimated with their respective posterior distributions, as demonstrated here in Table 4, Figures 4, 5 and 6. This is useful in interpreting several aspects of brain studies, from individual regions to RPs and network.
3. Extraction of region effects. A unique feature of the present approach is that it can infer region-level effects (in addition to RP effects, and the remaining additive contributions). In this manner, BML more accurately characterizes the contribution of a region through an integrative model that leverages the hierarchical effects at multiple levels. We propose that this property allows the assessment of region “importance” in a manner that is statistically more nuanced than those commonly used in graph theoretic analysis (such as “hubs” and “degrees”) and NBS. In addition, note that region-level effects at present cannot be obtained through the current alternative GLM-based methodologies such as NBS and FSLnets. The unique feature of statistical inference at the region level under BML is related to, but different from, some effort in the literature that associated the interrelationships among brain regions with the effects at those regions. For example, one study explored how the BOLD responses to tasks at a list of parcellated regions were related to the coherence matrix among those regions (Murphy et al., 2016). The two entities came from two distinct processes: the BOLD response at each region were estimated through typical time series regression at the individual subject level, while the coherence matrix of the regions was constructed between wavelet coefficients extracted from wavelet transform. In contrast, the two entities out of the BML approach for MBA are intertwined as the two sides of the same coin: as the formulas for region effect (9) and for RP (8) indicate, both effects at region and RP levels are assembled with the same components from one integrative model.
4. Integration and efficiency. BML builds an integrative platform and achieves high efficiency through sharing and pooling information across all entities involved in the system. Specifically, instead of modeling each RP separately as with the conventional GLM approaches, BML incorporates multiple testing as part of the model by assigning a prior distribution (e.g., Gaussian) among the ROIs (i.e., treating ROIs as random effects under the LME paradigm). In doing so, multiple testing is handled under the scaffold of the multilevel data structure by conservatively shrinking the original effect toward the center; that is, instead of leveraging cluster size or effect strength, BML leverages the commonality among ROIs. Compared to graph theoretic analysis in which the matrix data are transformed into a vector of topological features, BML for MBA maintains the integrity of the effect magnitudes at both region and RP levels.
5. Full reporting. The availability of full report through posterior distributions (e.g. Fig. 3) under BML provides rich and detailed information about each effect of interest, and avoids the need for thresholding under NHST and the resulting artificial dichotomization and results instabilities. We strongly suggest the practice of full results reporting through highlighting without, instead of the common practice of, hiding. Furthermore, full results reporting is more suited for reproducibility and meta analysis. For example, even if the evidence for an effect is at the level of 87% quantile interval, not high enough to reach a comfort zone of, for example, 95%, full results reporting would allow the evidence as a reference for future studies and for its inclusion in potential meta analysis. In contrast, the current practice of sweeping under the rug the results below the threshold is not only wasteful in data information (as well as resources, time and effort), but also detrimental for data reproducibility.
6. Flexibility. BML offers a flexible approach to dealing with double sidedness at the ROI and RP level. When prior information about the directionality of an effect is available on some, but not all, regions (e.g., from previous studies in the literature), one may face the issue of performing two one-tailed *t*-tests at the same time in a blindfold fashion due to the limitation of the massively univariate approach in the current practice (Chen et al., 2018b). The BML approach disentangles the complexity since the posterior inference for each ROI or RP can be made separately.
7. Validation. Model validation is a crucial facet of a Bayesian workflow framework. It is sometimes stated a model-or assumption-free approach is preferable to parametric methods, for example, with the argument that the P-value from permutation testing can be considered “exact.” However, it should be noted that “exactness” in this sense is a technical definition, requiring strict exchangeability and based on the assumption that the only possible values obtainable in an experiment were obtained. That is, the P-value’s “exactness” remains conditional on the current data. This emphasis on strict exactness under the NHST is questionable since the *P*-value itself is a random variable and the data are usually quite noisy; for instance, a repeated experiment under the same conditions would lead to a different “exact” *P*-value.

Furthermore, the capability of model checking (e.g., PPCs, Fig. 4) and validation (e.g., leave-one-out cross-validation, Table 2) under BML is unmatched from the conventional GLM framework (including permutation testing) in MBA. The determination coefficient *R*^2^ in a classic statistical model measures the proportion of the variance in the data that can be accounted for by the explanatory variable(s). However, it does not provide a well-balanced metric for model performance. For instance, a regression model fitted with high-order polynomials may behave well with the current data, but its predictability with a new dataset could fail severely. Therefore, a more reasonable approach to measure the predictive power of a model is to probe its applicability on a new dataset. However, instead of splitting between training and test data in machine learning, a much more efficient and sophisticated approach is to manipulate the data at hand through LOO-CV. It is an apparently widespread misconception that cross-validation seems to be only applicable to data mining and classifications. Instead, cross-validation should be adopted as an essential statistical technique, as demonstrated here in Table 2.

##### Limitations of BML in handling MBA

BML contains substantial modeling power and capabilities for MBA, but there are also a variety of challenges and risks. There are a few limitations with BML modeling. First, the computational cost could be high. As the number of regions *m* and the number of subjects *n* increase, the runtime can be hours, days, or even months. At present parallelization can only be achieved across chains, but not within each chain. However, currently there are some developments to take advantage of multi-threading (or MPI), which may realize within-chain parallelization in the future. Second, it might be problematic to make the assumption of effect decomposition with each RP effect *z_ijk_* as the additive effects of multiple components under BML. Admittedly, such additivity, as adopted in most statistical models, is bound to be vulnerable for potential assumption violations. However, we are compelled to cite the famous quote by G.E. Box, “all models are wrong, but some are useful.” Despite the vulnerability and the potential risk of poor fitting, we emphasize two aspects in the defense of our adoption of the BML framework: (a) the model quality can be directly assessed against the conventional approaches through various validation methods such as LOO-CV and PPCs, as illustrated in Fig. 3, and (b) any problems under the current model serve as a starting point for cumulative and recursive modeling improvement process in the future. Third, one concern is that the exchangeability requirement of BML assumes that no differential information is available across the ROIs in the model. Exchangeability captures symmetry among the ROIs in a sense that does not require independence. That is, an independent and identically distributed set of entities (e.g., ROIs) is exchangeable, but not vice versa. However, every exchangeable set of entities (e.g., ROIs) is identically distributed (Gelman et al., 2014). Under some circumstances, ROIs can be expected to share some information and not fully independent, especially when they are anatomically contiguous or more functionally related than the other ROIs (e.g., homologous regions in opposite hemisphere). However, the exchangeability is an epistemological, neither physical nor ontological, assumption that renders a convenient approximation of a prior distribution by a mixture of i.i.d. distributions (de Finetti’s theorem, Gelman et al., 2014). The presence of temporal correlation in time series regression may cause the underestimation of variances because the conventional statistics heavily relies on the concept of degrees of freedom. In contrast, Bayesian inferences build on posterior distributions without invoking the degrees of freedom, and the violation of exchangeability usually leads to negligible effect on the final shape of posterior distributions except for the precise sequence in which the posterior draws occur (McElreath, 2016).

#### Conclusion

The proposed Bayesian multilevel (BML) modeling framework can be applied generally to any matrix-based analysis (MBA). Through decomposing the effect at each element of the data matrix into multiple components such as row, column, subject as well as their interactions, we have built an LME and its BML counterpart to more accurately capture the data structure and the intertwining relationships. More importantly, as a principled compromise between local and global effects through partial poolig, the BML framework allows the investigator to more efficiently make statistical inferences at both region and region pair levels under one unified model, and avoids the penalty for multiple testing involved in the conventional GLM approach in which each region pair is treated as an unrelated entity. The inferences at each region under BML offer a unique effect characterization; furthermore, full results reporting with BML avoids unnecessary loss of information and waste of effort and research resources that plague the methods associated with sharp thresholding.

## Acknowledgments

The research and writing of the paper were supported (GC, PAT, and RWC) by the NIMH and NINDS Intramural Research Programs (ZICMH002888) of the NIH/HHS, USA. LP’s research is supported in part by the National Institute of Mental Health (R01 MH071589 and R01 MH112517). We thank Vitria Adisetiyo for her help and are indebted to the Stan development team for their technical support. The computations were performed and the figures were generated with the R language for statistical computing (R Core Team, 2018). This work utilized the computational resources of the NIH HPC Biowulf cluster.(http://hpc.nih.gov).

## Footnotes

1 Several variations of correlation exist such as Pearson correlation between the time courses of two components from ndependent (or principal) component analysis, partial correlation and spectral coherence.

2 The effect decomposition of the BML model remains the same as its LME counterpart. The different model expression he e is adopted to emphasize the fact that the outcome under BML is conditional on the parameters and priors.

3 In theory, the multi-membership scheme can also be, but has not been yet, implemented under conventional LME.

4 See https://en.wikipedia.org/wiki/Folded-t_and_half-t_distributions for the density *p*(*v μ*, *σ*^2^) of folded non-stand rdized *t*-distribution, where the parameters *v μ*, and *σ*^2^ are the degrees of freedom, mean, and variance.

5 The LKJ prior (Lewandowski, Kurowicka, and Joe, 2009) is a distribution over symmetric positive-definite matrices w th the diagonals of 1s.

6 The posterior probability of the effect being negative at a region is *p*_=1 −*p*_+._

7 Such a trial-and-error methodology is even encouraged in the NBS manual (most recent version, v1.2, at the time of writing; https://www.nitrc.org/projects/nbs/): “We recommend experimenting with a range of thresholds. Sensitivity to the test statistic threshold may reveal useful information about the nature of the effect.”

